# Intracellular pH dynamics promotes zebrafish larval tail regeneration

**DOI:** 10.64898/2026.05.14.724960

**Authors:** Cambria Chou-Freed, Christopher K Prinz, Anush Margaryan, Julie A Theriot, Daniel E Wagner, Diane L Barber

**Affiliations:** Department of Cell and Tissue Biology, University of California San Francisco, San Francisco, CA 94143, USA; Department of Biology, University of Washington, Seattle, WA 98195, USA; Howard Hughes Medical Institute, University of Washington, Seattle, WA 98195, USA; Developmental & Stem Cell Biology Program, University of California San Francisco, San Francisco, CA 94143, USA; Department of Obstetrics, Gynecology and Reproductive Science, Center for Reproductive Sciences, University of California San Francisco, San Francisco, CA 94143, USA; Eli and Edythe Broad Center for Regeneration Medicine and Stem Cell Research, University of California San Francisco, San Francisco, CA 94143, USA

## Abstract

Intracellular pH (pHi) dynamics regulates numerous cell behaviors, including migration and proliferation. While these functions are well-established in cell lines, the role of pHi changes *in vivo* is less well understood. We generated a transgenic zebrafish line expressing a fluorescent ratiometric pHi biosensor and identified functional changes in pHi during zebrafish larval tail regeneration. We found that tail amputation led to a transient decrease in pHi, followed by a prolonged increase in pHi above pre-amputation values. Moreover, we showed that pharmacologically inhibiting Na^+^/H^+^ exchanger (NHE) activity or decreasing extracellular pH attenuated the post-amputation increase in pHi, reduced subsequent cell proliferation, and impaired tail regeneration. We further found that inhibiting NHE activity post-amputation led to elevated inflammation, disrupted myeloid cell behavior, decreased reactive oxygen species, and increased glycogen synthase kinase-3 (GSK3) activity. Finally, we showed that the regeneration defects in larvae with disrupted pHi were partially rescued by the GSK3 inhibitor BIO. Our data reveal a previously unrecognized role for pHi dynamics in coordinating tissue behaviors *in vivo* and enabling zebrafish larval tail regeneration.

## INTRODUCTION

Intracellular pH (pHi) is highly dynamic, playing a critical role in regulating cell behaviors across species, from bacteria to humans. Coordinated by plasma membrane ion transporters such as Na^+^/H^+^ exchangers (NHEs) (Orlowski and Grinstein, 2011) and HCO_3_^-^ transporters (Romero et al., 2013), an increase in pHi is required for cell migration (Denker and Barber, 2002; Frantz et al., 2007; Patel and Barber, 2005; Stock, 2024), proliferation (Flinck et al., 2018; Putney and Barber, 2003; Spear and White, 2023), metabolic reprogramming (Kazyken et al., 2023; Man et al., 2022; Manoli et al., 2021), and cell fate decisions (Liu et al., 2023; Ulmschneider et al., 2016). While most reports on pHi dynamics use clonal cells, a limited number of *in vivo* studies have revealed how increased pHi is necessary for tail bud elongation in chick embryos (Oginuma et al., 2020), melanocyte maturation in zebrafish embryos (Raja et al., 2020), ovary development (Benitez et al., 2019; Ulmschneider et al., 2016) and tumorigenesis (Grillo-Hill et al., 2015) in *Drosophila*, and defecation (Pfeiffer et al., 2008) and lifespan (Nehrke, 2003) in *C. elegans*. However, determining pHi *in vivo* remains challenging, largely due to limitations in methods for quantifying and perturbing pHi in live animals. To address this challenge, we generated a transgenic zebrafish line expressing a fluorescent ratiometric pHi biosensor, mKate2-SEpHluorin (Miesenböck et al., 1998; Tanida et al., 2014), which enabled us to quantify changes in pHi during complex tissue behaviors *in vivo*. Using this new line along with calibration methods, we quantified pHi during zebrafish larval tail regeneration, a process similar to adult tail regeneration (Kawakami et al., 2004).

Zebrafish larval tail regeneration involves many of the aforementioned cell behaviors that are known to depend on increased pHi in other contexts. The process begins with the wound healing phase, in which damage-induced signals such as reactive oxygen species (ROS) (Niethammer et al., 2009) coordinate the contraction and migration of epidermal cells to close the wound (Ding et al., 2025; Gault et al., 2014; Kawakami et al., 2004), as well as the recruitment of migrating neutrophils and macrophages (Li et al., 2012; Nguyen-Chi et al., 2017; Sipka et al., 2021). While these myeloid cells initially serve crucial pro-inflammatory functions, neutrophils later leave the wound site through reverse migration (Mathias et al., 2006; Tauzin et al., 2014; Yoo and Huttenlocher, 2011), and macrophages switch to an anti-inflammatory, pro-regenerative phenotype (Hasegawa et al., 2017; Nguyen-Chi et al., 2017). During the subsequent proliferative phase of regeneration, a cascade of signaling pathways induces cell proliferation and specification of new cell types, including a layer of specialized epithelial cells called the wound epithelium, as well as the blastema, a population of mesenchymal cells that gives rise to regenerated tissue (Kawakami et al., 2004; Mathew et al., 2009; Romero et al., 2018).

We hypothesized that pHi dynamics might play a role in one or more of these processes during zebrafish larval tail regeneration. In support of this hypothesis, a previous study found that tail amputation in *Xenopus laevis* leads to a rapid decrease in pHi that recovers by 1 hour post-amputation (hpa) (Matlashov et al., 2015)—although the functional role of these pHi changes in the regeneration process remains unclear. Using our *Tg(ef1α:mKate2-SEpHluorin)* pHi biosensor line, we found that zebrafish larval tail amputation led to an initial transient decrease in pHi, followed by a prolonged increase in pHi above pre-amputation levels. We further found that attenuating the post-amputation increase in pHi by pharmacologically inhibiting NHE activity or by decreasing extracellular pH (pHe) led to reduced cell proliferation and impaired tail regeneration. We showed that inhibiting NHE activity post-amputation was associated with elevated inflammation, disrupted myeloid cell behavior, reduced ROS production, and increased glycogen synthase kinase-3 (GSK3) activity. Finally, we found that the regeneration and proliferation defects in larvae with disrupted pHi were partially rescued by co-treatment with the GSK3 inhibitor BIO. Collectively, our data reveal previously unrecognized roles for pHi dynamics in promoting zebrafish larval tail regeneration.

## RESULTS

### A transgenic biosensor reveals pHi dynamics following zebrafish larval tail amputation

To quantify pHi in zebrafish, we generated a transgenic line expressing a nucleocytoplasmic ratiometric pHi biosensor, mKate2-SEpHluorin, (Tanida et al., 2014) driven by a ubiquitous ef1*α* driver (Fig. 1A). Superecliptic pHluorin (SEpHluorin) (Miesenböck et al., 1998), with a pKa of ∼7.1 (Sankaranarayanan et al., 2000), increases fluorescence intensity linearly with increased pH within a physiologically relevant range, while mKate2, with a pKa of ∼5.4 (Shcherbo et al., 2009), is largely insensitive to changes in pH. Thus, measured ratios of SEpHluorin to mKate2 fluorescence intensity can be used as an indicator of pHi normalized to biosensor expression.

**Fig. 1.**
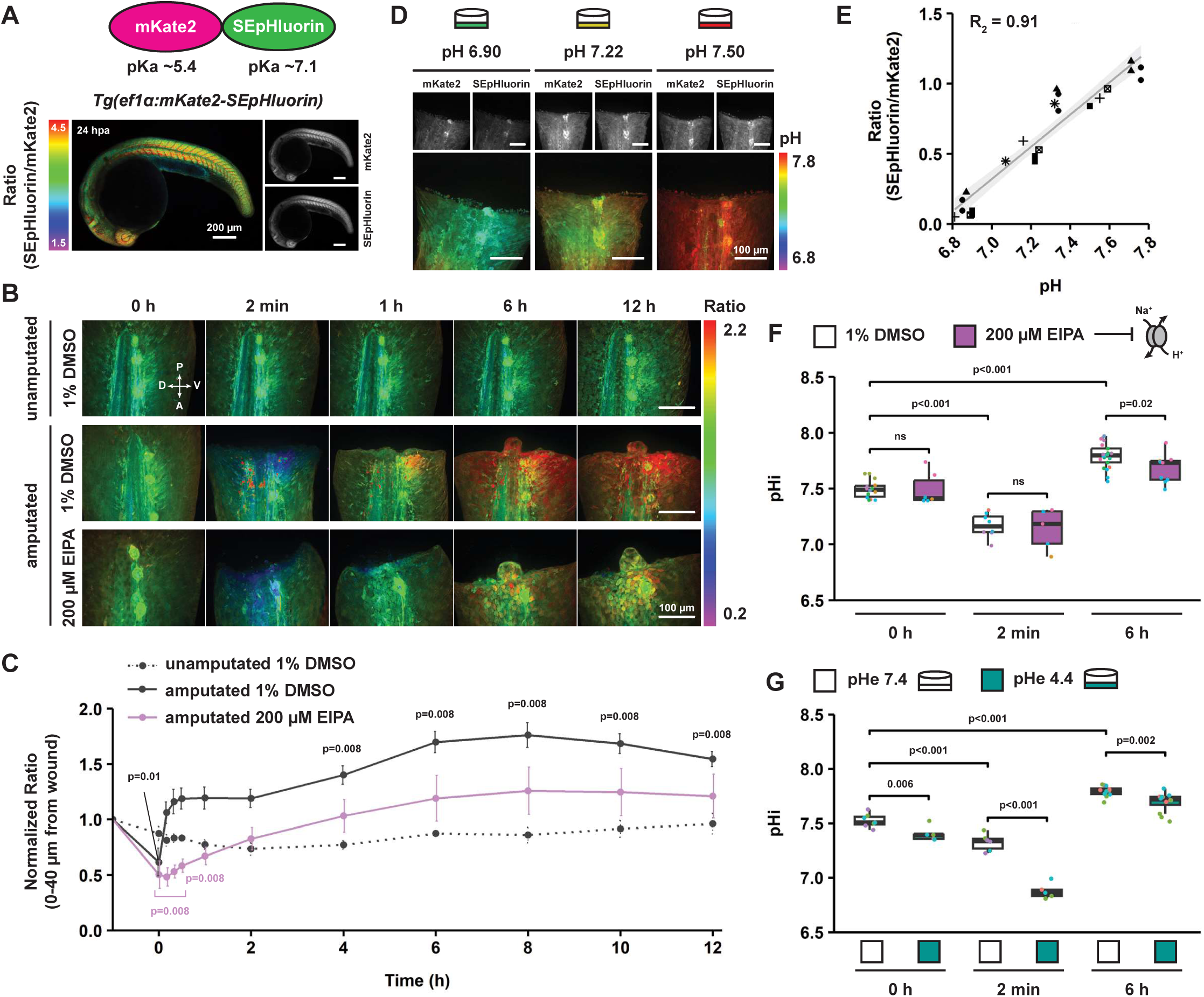
Increased pHi following tail amputation is attenuated by the NHE inhibitor EIPA or low pHe. (A) Schematic illustrating pHi biosensor mKate2-SEpHluorin and representative maximum projection images of a *Tg(ef1α:mKate2-SEpHluorin)* embryo at 24 hpa. Scale bars: 200 µm. (B) Representative maximum projection images of SEpHluorin/mKate2 ratios in unamputated and amputated *Tg(ef1α:mKate2-SEpHluorin)* larval tails at 3 dpf over 12 h. Scale bars: 100 µm. P – posterior, A – anterior, D – dorsal, V – ventral. (C) Time course of SEpHluorin/mKate2 ratios measured in unamputated and amputated *Tg(ef1α:mKate2-SEpHluorin)* larval tails (0-40 µm from wound) normalized to t=0 timepoint for each fish. Error bars show SEM. n=5,5,3. p-values from Kruskal-Wallis (n.s. for unamputated group; p<0.001 for amputated groups) and Wilcoxon signed rank tests (mu=1). (D) Representative maximum projection images of amputated *Tg(ef1α:mKate2-SEpHluorin)* larval tails at 3 dpf incubated for 20 min in nigericin calibration buffers. (E) SEpHluorin/mKate2 ratios measured in amputated *Tg(ef1α:mKate2-SEpHluorin)* larval tails (0-40 µm from wound) incubated for 20 min in nigericin calibration buffers plotted vs. pH of buffers. Dots represent fish. Dot shapes represent experimental days. n=28. (F,G) Calibrated pHi values measured in unamputated (0 h) and amputated *Tg(ef1α:mKate2-SEpHluorin)* larval tails (0-40 µm from wound) at indicated timepoints. Dots represent fish. Dot colors represent experimental days. Boxplots show median and interquartile range. n=6-21 (F) or n=5-12 (G) per group. p-values from two-way ANOVA and pairwise t-tests with Benjamini-Hochberg correction.

Using this *Tg(ef1α:mKate2-SEpHluorin)* line, we quantified pHi changes in larval tail tissue following amputation at the posterior end of the pigment gap (as described in Romero et al., (2018)). This “tail regeneration” paradigm differs from “fin fold regeneration” and represents a more complex regenerative challenge involving the partial removal of neural tube, notochord, muscle, and other cell types in addition fin fold epithelium and mesenchyme. We conducted *in vivo* time lapse imaging of unamputated and amputated *Tg(ef1α:mKate2-SEpHluorin)* larvae at 3 days post-fertilization (dpf) and quantified SEpHluorin/mKate2 ratios over 12 hours. In unamputated tails, SEpHluorin/mKate2 ratios remained constant (Fig. 1B,C; Fig. S1A). By contrast, amputated tails displayed a decrease in ratios at 2 minutes post-amputation (mpa) that recovered by 10 mpa, followed by an increase in ratios above pre-amputation values beginning at 4 hpa and reaching a maximum at 8 hpa (Fig. 1B,C; Fig. S1A). These ratio changes occurred most prominently in cells within 40 µm of the amputation plane but were also observed up to 120 µm from the wound site (Fig. S2).

We quantified pHi values from SEpHluorin/mKate2 ratios using standard calibration methods with the K^+^/H^+^ ionophore nigericin (see Materials and Methods). SEpHluorin/mKate2 ratios were linearly correlated with nigericin buffer pH within a pH range of 6.8 to 7.8 (R² = 0.91) (Fig. 1D,E), largely due to changes in SEpHluorin (R² = 0.78) rather than mKate2 (R² = 0.40) fluorescence intensity (Fig. S1B,C). Using internal calibration curves collected for each biological replicate, we determined that pre-amputation tails displayed a mean pHi of 7.49, which decreased to 7.17 at 2 mpa and increased to 7.79 at 6 hpa (Fig. 1F). These data indicate that larval tail amputation triggers widespread changes in pHi over 12 hpa.

### Increased pHi post-amputation is attenuated by the NHE inhibitor EIPA and low pHe

To determine the functional significance of pHi dynamics during tail regeneration, we sought approaches to perturb pHi in amputated tails. We first tested 5-(N-ethyl-N-isopropyl)-amiloride (EIPA), a pharmacological inhibitor of NHE activity. We incubated larvae in 200 µM EIPA for 12 hpa and found that SEpHluorin/mKate2 ratios decreased at 2 mpa similarly to 1% DMSO vehicle-treated controls (Fig. 1B,C; Fig. S1A). However, ratios recovered more slowly with EIPA (by 2 hpa) and failed to subsequently increase beyond pre-amputation levels (Fig. 1B,C; Fig. S1A). Standard calibration of these ratios confirmed that pHi was reduced with EIPA relative to DMSO at 6 hpa (mean value of 7.68 in EIPA), with no differences at 2 mpa or pre-amputation (Fig. 1F).

As a second approach to experimentally change pHi, we incubated larvae in buffered isotonic media of different pH values for up to 6 hpa. We found that incubating larvae in a neutral pHe of 7.4 resulted in similar pHi dynamics as in DMSO-treated controls, with mean pHi values of 7.53 pre-amputation, 7.32 at 2 mpa, and 7.79 at 6 hpa (Fig. 1G). By contrast, incubating larvae in a low pHe of 4.4 led to a decrease in pHi (mean values of 7.40 pre-amputation, 6.88 at 2 mpa, and 7.69 at 6 hpa) (Fig. 1G). Thus, exposure to either EIPA or low pHe leads to reduced pHi in amputated larval tails in the first 12 hpa.

### Decreasing pHi with EIPA or low pHe impairs tail regeneration

To determine the role of pHi dynamics during tail regeneration, we asked whether sustained exposure to EIPA or low pHe affects tail regrowth. We found that EIPA treatment decreased area of regrowth relative to DMSO-treated controls at 3 dpa in a dose-dependent manner, with 175 and 200 µM EIPA having a significant effect (50% reduction in area with 200 µM EIPA) (Fig. 2A,B). Additionally, we found that a low pHe of 4.4, but not a high pHe of 8.9, decreased area of regrowth by 23% compared with a neutral pHe of 7.4 (Fig. 2C,D). By contrast, incubating unamputated larvae from 3 to 6 dpf in 200 µM EIPA or pHe 4.4 had no effect on larval survival or tail growth (Fig. S3). These data indicate that limiting the increase in pHi post-amputation impairs larval tail regeneration.

**Fig. 2.**
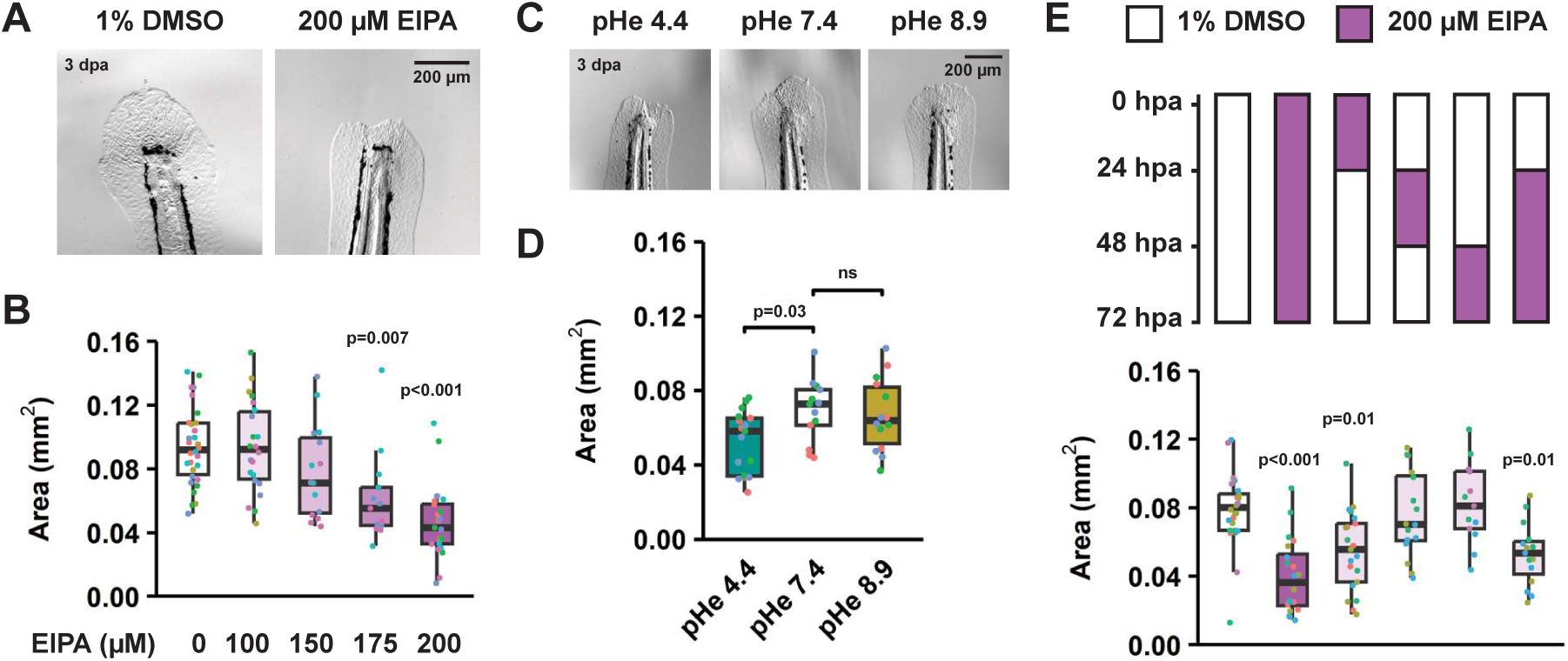
Post-amputation exposure to EIPA or low pHe impairs tail regeneration. (A,B) Representative images (A) and area of regrowth (B) of wildtype larval tails amputated in increasing concentrations of EIPA at 3 dpa. n=13-30 per group. p-values from Kruskal-Wallis (p<0.001) and Dunn’s tests. (C,D) Representative images (C) and area of regrowth (D) of wildtype larval tails amputated in indicated pHe at 3 dpa. n=13-17 per group. p-values from one-way ANOVA (p=0.02) and Dunnett’s tests vs. pHe 7.4. (E) Area of regrowth measured in amputated wildtype larval tails at 3 dpa after treatment with 200 µM EIPA during different time windows. n=13-28 per group. p-values from Kruskal-Wallis (p<0.001) and Dunn’s tests. (B,D,E) Dots represent fish. Dot colors represent experimental days. Boxplots show median and interquartile range.

To define the temporal dependence of pHi changes during the regeneration process, we quantified area of regrowth after treating larvae with 200 µM EIPA during explicit exposure windows. EIPA treatment for 24 hours on day 1 post-amputation, but not on day 2 or 3, was sufficient to decrease area of regrowth by 30% (Fig. 2E). EIPA treatment for 48 hours on days 2 to 3 led to a similar regrowth deficit of 33% (Fig. 2E). These results indicate an early window of sensitivity over the first 24 hours post-amputation that coincides with the timing of post-amputation changes in pHi described above, followed by a late window of more moderate sensitivity over the subsequent 48 hours. These time windows roughly correspond to the wound healing and proliferative phases of regeneration, respectively, suggesting a role for pHi dynamics during both processes.

Based on our findings with EIPA, we asked whether genetic disruption of NHEs also leads to impaired tail regeneration. In mammals, NHE1 is the ubiquitously expressed, resident plasma membrane isoform that commonly regulates pHi (Orlowski and Grinstein, 2011). Zebrafish have 10 NHE family members, with NHE1a (encoded by the *slc9a1a* gene) having the greatest sequence similarity to human NHE1 (Fig. S4A). Because few zebrafish NHE isoforms have been functionally characterized, we conducted a targeted CRISPR/Cas9 screen of four NHE genes encoding predicted plasma membrane isoforms: *slc9a1a*, *slc9a1b*, *slc9a2*, and *slc9a8* (Orlowski and Grinstein, 2011). We measured tail regrowth at 3 dpa in F_0_ crispant larvae and found that mosaic targeting of each of the four genes decreased area of regrowth relative to controls injected with a scrambled sgRNA (Fig. S4B). By contrast, unamputated crispant larvae targeted for each of the four genes showed no differences in tail growth (Fig. S4C). Analysis of homozygous null larvae resulting from an incross between stable heterozygous carriers of premature stop codons revealed no differences in area of regrowth at 3 dpa relative to wildtype or heterozygous siblings (Fig. S4D-G), a result possibly explained by genetic compensation triggered by mutant mRNA (El-Brolosy et al., 2019).

### Bulk RNA-sequencing reveals early and late transcriptional responses to EIPA treatment during tail regeneration

Because disrupting pHi during either the wound healing or the proliferative phase impaired tail regeneration, we hypothesized that pHi dynamics contributes to tail regeneration via multiple mechanisms. To begin identifying these mechanisms, we conducted bulk RNA-sequencing (RNA-seq) of DMSO- and EIPA-treated larvae at both an early timepoint (6 hpa) during the wound healing phase and a late timepoint (48 hpa) during the proliferative phase.

At the early 6 hpa timepoint, we sequenced tail tissue from amputated DMSO-and EIPA-treated larvae (Fig. S5A). Principal component analysis revealed reproducible transcriptomic differences between DMSO- and EIPA-treated samples (Fig. S5B). Relative to DMSO-treated controls, EIPA treatment resulted in the upregulation of 60 genes and downregulation of 53 genes (adjusted p-value < 0.05) (Supp Table 1).

**Table 1.**
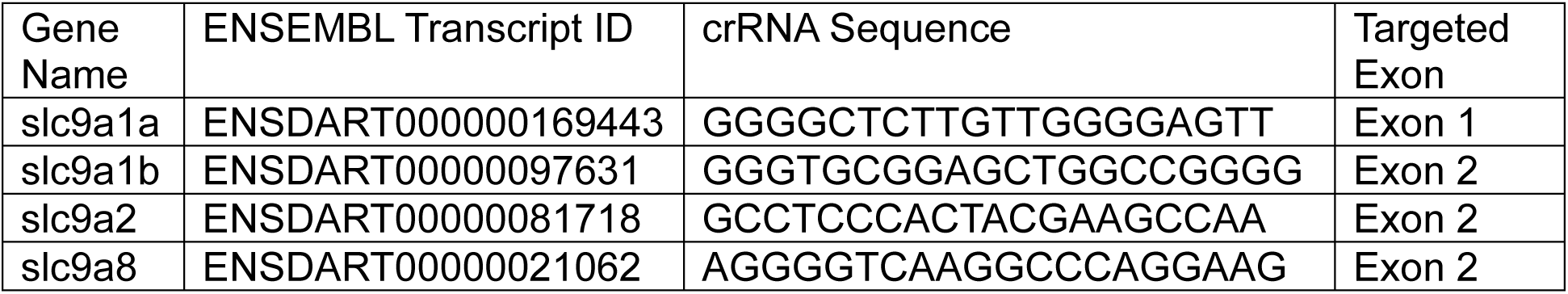
CRISPR/Cas9 crRNAs.

Among the genes downregulated following EIPA treatment, gene ontology (GO) analysis revealed cholesterol and isoprenoid biosynthesis as top enriched biological processes (Fig. S5C; Supp Table 2). Specifically, EIPA treatment led to the downregulation of five genes encoding enzymes within the cholesterol biosynthetic pathway, including *hmgcs1*, *hmgcra*, *fdps*, *sqlea*, and *cyp51* (Fig. S5D,E), suggesting that cholesterol biosynthesis might play a role in the regeneration process. We therefore targeted Hmgcr with the pharmacological inhibitor atorvastatin and found that 10 µM atorvastatin decreased area of regrowth by 29% relative to DMSO-treated controls at 3 dpa (Fig. S5F,G), with no effect on unamputated larval survival or tail growth (Fig. S5H-J). These data suggest a potential role for cholesterol biosynthesis during zebrafish larval tail regeneration.

**Table 2.**
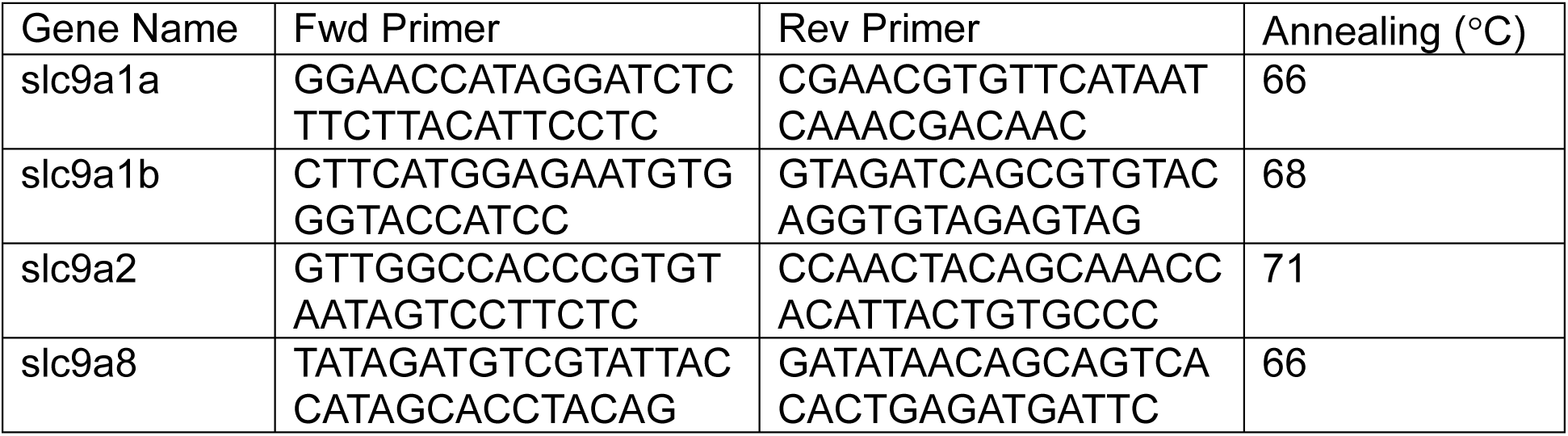
PCR Primers for T7E1 Assays.

To determine the consequences of EIPA treatment during the proliferative phase of regeneration, we sequenced tail tissue from amputated and unamputated larvae treated for 48 hours with either DMSO or EIPA (Fig. 3A). In parallel, to reveal the specific consequences of early exposure to EIPA, we also sequenced tail tissue from amputated larvae treated for 24 hours with EIPA followed by DMSO “washout” of the drug from 24 to 48 hpa (Fig. 3A). Principal component analysis of all samples revealed transcriptomic effects of both EIPA treatment and regenerative challenge, with the amputated EIPA washout samples co-clustering with the amputated DMSO controls (Fig. 3B). This result suggests that late EIPA treatment (after 24 hpa) exerts distinct and strong effects during the proliferative phase.

**Fig. 3.**
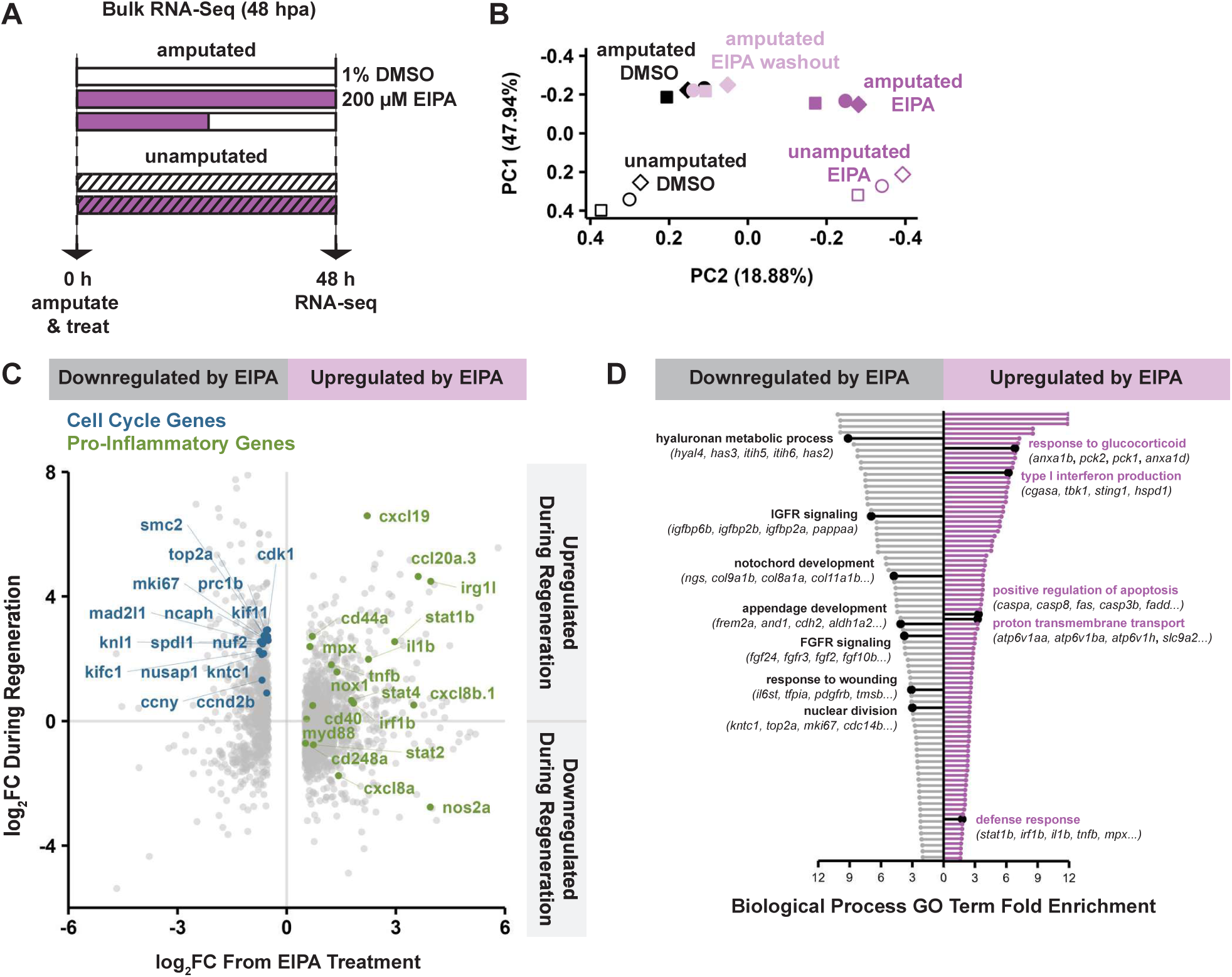
Bulk RNA-seq at 48 hpa reveals transcriptional consequences of EIPA treatment during the proliferative phase. (A) Schematic of bulk RNA-seq conditions at 48 hpa. (B) Principal component analysis (PCA) plot of bulk RNA-seq results at 48 hpa. Dots represent samples. Dot shapes represent replicates. (C) Scatterplot of differentially expressed genes (DEGs) between amputated EIPA- and DMSO-treated larvae at 48 hpa. x-axis shows log_2_FC (FC – fold change) from EIPA treatment (amputated: EIPA vs. DMSO comparison). y-axis shows log_2_FC from regenerative challenge (DMSO: amputated vs. unamputated comparison). (D) Gene ontology (GO) terms enriched among DEGs between amputated EIPA- and DMSO-treated larvae at 48 hpa. IGFR – insulin-like growth factor receptor, FGFR – fibroblast growth factor receptor.

Identifying differentially expressed genes (DEGs) between conditions further revealed transcriptional changes linked to EIPA treatment, regenerative challenge, or combined effects. We first identified general signatures of EIPA treatment by comparing EIPA- and DMSO-treated samples from both amputated and unamputated larvae. In amputated larvae, EIPA treatment resulted in the differential expression (adjusted p-value < 0.05 and |log_2_ fold change| > 0.5) of 2171 genes (1421 upregulated; 750 downregulated) (Fig. 3C; Fig. S6A; Supp Table 1). A large proportion of these DEGs (63%; 1369 genes) also responded in unamputated larvae (Fig. S6B; Supp Table 1), revealing a specific signature of EIPA treatment irrespective of regenerative challenge. Meanwhile, EIPA washout treatment resulted in the differential expression of 51 genes (27 upregulated; 24 downregulated), of which a large proportion (71%; 36 genes) responded similarly to full-length EIPA treatment (Fig. S6B; Supp Table 1). This result reveals a highly specific set of transcriptional changes shared between early and full-length EIPA treatment conditions.

Separate differential gene expression tests comparing amputated and unamputated DMSO-treated controls revealed a baseline set of DEGs that were globally upregulated (n = 4135) or downregulated (n = 4269) during the proliferative phase of regeneration (Fig. S6A; Supp Table 1). We compared this process in EIPA-treated larvae and found that EIPA treatment prevented the response of many of these regeneration-specific genes: 43% of DEGs normally upregulated during the proliferative phase (n = 1830) and 38% (n = 1558) of DEGs normally downregulated failed to respond to regenerative challenge in EIPA-treated larvae (Fig. S6C; Supp Table 1). These analyses indicate that EIPA treatment has reproducible transcriptional effects that intersect with the transcriptional signature of the proliferative phase.

Among the genes upregulated following full-length EIPA treatment in amputated larvae, GO analysis revealed the enrichment of genes associated with proton transmembrane transport, including the NHE gene *slc9a2* and multiple V-ATPase subunit genes (Fig. 3D; Supp Table 2). By contrast, genes downregulated following full-length EIPA treatment in amputated larvae were enriched for genes associated with response to wounding, appendage development, and notochord development, as well as multiple signaling pathways known to be critical for zebrafish larval tail regeneration, including fibroblast growth factor (FGF) (Kawakami et al., 2004; Mathew et al., 2009; Romero et al., 2018), insulin-like growth factor (IGF) (Lewis et al., 2023), and hyaluronan (Ouyang et al., 2017) (Fig. 3D; Supp Table 2). GO analysis further revealed many additional biological processes impacted by EIPA treatment in amputated larvae, including nuclear division, apoptosis, and inflammatory processes (Fig. 3D; Supp Table 2), supporting our hypothesis that disrupting pHi dynamics impairs tail regeneration through multiple mechanisms. We chose to focus on two of these suggested mechanisms, proliferation and inflammation.

### Decreasing pHi with EIPA or low pHe leads to cell proliferation defects during tail regeneration

Our RNA-seq results at 48 hpa indicated that EIPA treatment in amputated but not unamputated larvae led to the downregulation of many cell cycle progression genes normally upregulated during regeneration, including the cyclin *ccny*, the cyclin-dependent kinase *cdk1*, the topoisomerase *top2a*, and the mitotic marker *mki67* (Fig. 3C,D; Fig. S6D; Supp Table 2). This result prompted us to test whether perturbing pHi post-amputation leads to impaired proliferation. We first immunolabeled for phosphorylated histone H3 (phospho-H3) and found an increased number of phospho-H3+ nuclei in DMSO-treated larvae at 48 hpa compared with unamputated larvae at the equivalent timepoint (Fig. 4A,B; Fig. S7A). EIPA treatment had no effect on the number of phospho-H3+ nuclei in unamputated larvae (Fig. 4A,B; Fig. S7A). However, in amputated larvae, EIPA treatment led to a reduced number of phospho-H3+ nuclei relative to DMSO-treated controls (Fig. 4A,B; Fig. S7A), a result consistent with previous findings that EIPA inhibits proliferation of clonal cells (Flinck et al., 2018; Putney and Barber, 2003). We confirmed this result by EdU labeling, which revealed fewer EdU+ nuclei with EIPA compared with DMSO at 48 hpa (Fig. S7C,D). We also found that incubating in low pHe led to a decreased number of phospho-H3+ nuclei at 48 hpa compared with neutral pHe (Fig. 4C,D; Fig. S7B). Together, these data suggest that limiting the increase in pHi post-amputation with either EIPA or low pHe leads to reduced cell proliferation during tail regeneration.

**Fig. 4.**
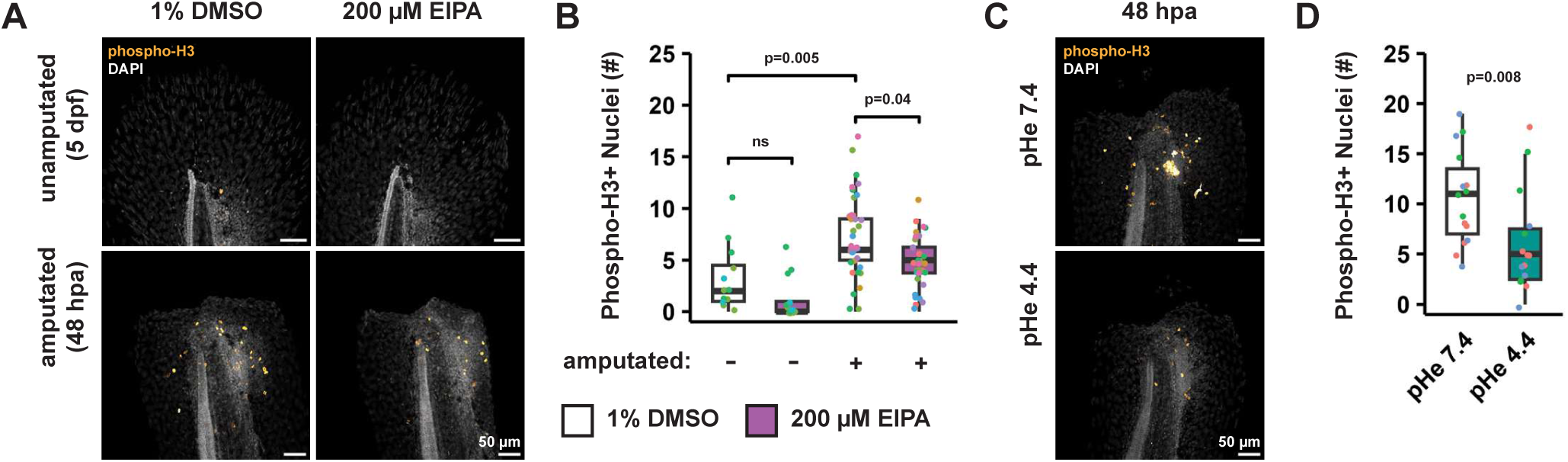
Post-amputation exposure to EIPA or low pHe leads to reduced cell proliferation. (A,B) Representative maximum projection images (A) and quantifications (B) of phospho-H3 immunolabeling in unamputated and amputated wildtype larval tails in 1% DMSO or 200 µM EIPA. n=12,13 (unamputated groups) or n=36,32 (amputated groups). p-values from Kruskal-Wallis (p<0.001) and Dunn’s tests. (C,D) Representative maximum projection images (C) and quantifications (D) of phospho-H3 immunolabeling in amputated wildtype larval tails in indicated pHe at 48 hpa. n=15 per group. p-value from Wilcoxon rank sum test. (B,D) Dots represent fish. Dot colors represent experimental days. Boxplots show median and interquartile range.

Our RNA-seq results at 48 hpa also indicated that EIPA treatment led to the upregulation of several pro-apoptotic genes, including caspases *caspa*, *casp8*, and *casp3b*, as well as the cell death receptor *fas* and its downstream signaling effector *fadd* (Fig. 3D; Supp Table 2). We therefore used TUNEL labeling, an indicator of fragmented DNA, to determine whether disrupting pHi promotes cell death. We found that EIPA treatment led to an increased number of TUNEL foci relative to DMSO-treated controls both at 48 hpa and in unamputated larvae at the equivalent timepoint (Fig. S8A,B). By contrast, the number of TUNEL foci was unchanged in low pHe compared with neutral pHe at 48 hpa (Fig. S8C,D). These results suggest that EIPA treatment may cause cell death independently of its effects on pHi, consistent with a previous report that EIPA triggers apoptosis in human cancer cells independent of NHE1 (Rolver et al., 2020), although further data would be required to confirm this possibility.

We next asked whether EIPA treatment leads to impaired specification of wound epithelium or blastema cells, the regenerative cells that initiate and sustain the proliferative response. Our RNA-seq data at 48 hpa indicated that EIPA treatment in amputated larvae resulted in a modest downregulation of both the wound epithelium marker *dlx5a* (log_2_FC -0.57) (Kawakami et al., 2004; Romero et al., 2018) and the blastema marker *aldh1a2* (log_2_FC -0.50) (Kawakami et al., 2004; Mathew et al., 2009; Romero et al., 2018) (Fig. S9A,B). However, quantification of *dlx5a* and *aldh1a2* expression by RNA fluorescence *in situ* hybridization (RNA-FISH) revealed increased *dlx5a* expression at the wound edge and unaltered levels of *aldh1a2* expression with EIPA relative to DMSO at 48 hpa (Fig. S9C-E). Notwithstanding these differences, both methods confirmed that *dlx5a* and *aldh1a2* were expressed to some degree in amputated larvae treated with EIPA, suggesting that EIPA does not fully impair specification of the wound epithelium or blastema. In summary, our findings suggest that decreasing pHi with EIPA or low pHe leads to defects in cell proliferation, which likely explain the impaired tail regeneration that we observed (Fig. 2A-D).

### EIPA treatment alters the inflammatory response and myeloid cell behavior during tail regeneration

A salient finding from our RNA-seq data at 48 hpa was that EIPA treatment in both amputated and unamputated larvae led to the upregulation of multiple immune pathway genes associated with defense response, type I interferon production (also upregulated at 6 hpa), and response to glucocorticoid (Fig. 3C,D; Supp Table 2). Most of these genes encode mediators of inflammation, including STAT transcription factors (*stat1b*, *stat2*, *stat4*), members of the STING1 network (*cgasa*, *sting1*, *myd88*), interferon signaling components (*irf1b*, *irf7*, *tbk1*, *trim25*), and inflammatory cytokines (*tnfb*, *cxcl19*, *ccl25b*, *il1b,* among others) (Fig. 3D; Fig. S6D). We confirmed increased expression of the pro-inflammatory cytokine *il1b* (interleukin-1β) by RNA-FISH in EIPA compared with DMSO at 48 hpa and in unamputated larvae at the equivalent timepoint (Fig. 5A,B). These data suggest that EIPA treatment leads to increased inflammation—a process which, though initially required for zebrafish larval tail regeneration, can be deleterious to regeneration in excess (Hasegawa et al., 2017).

**Fig. 5.**
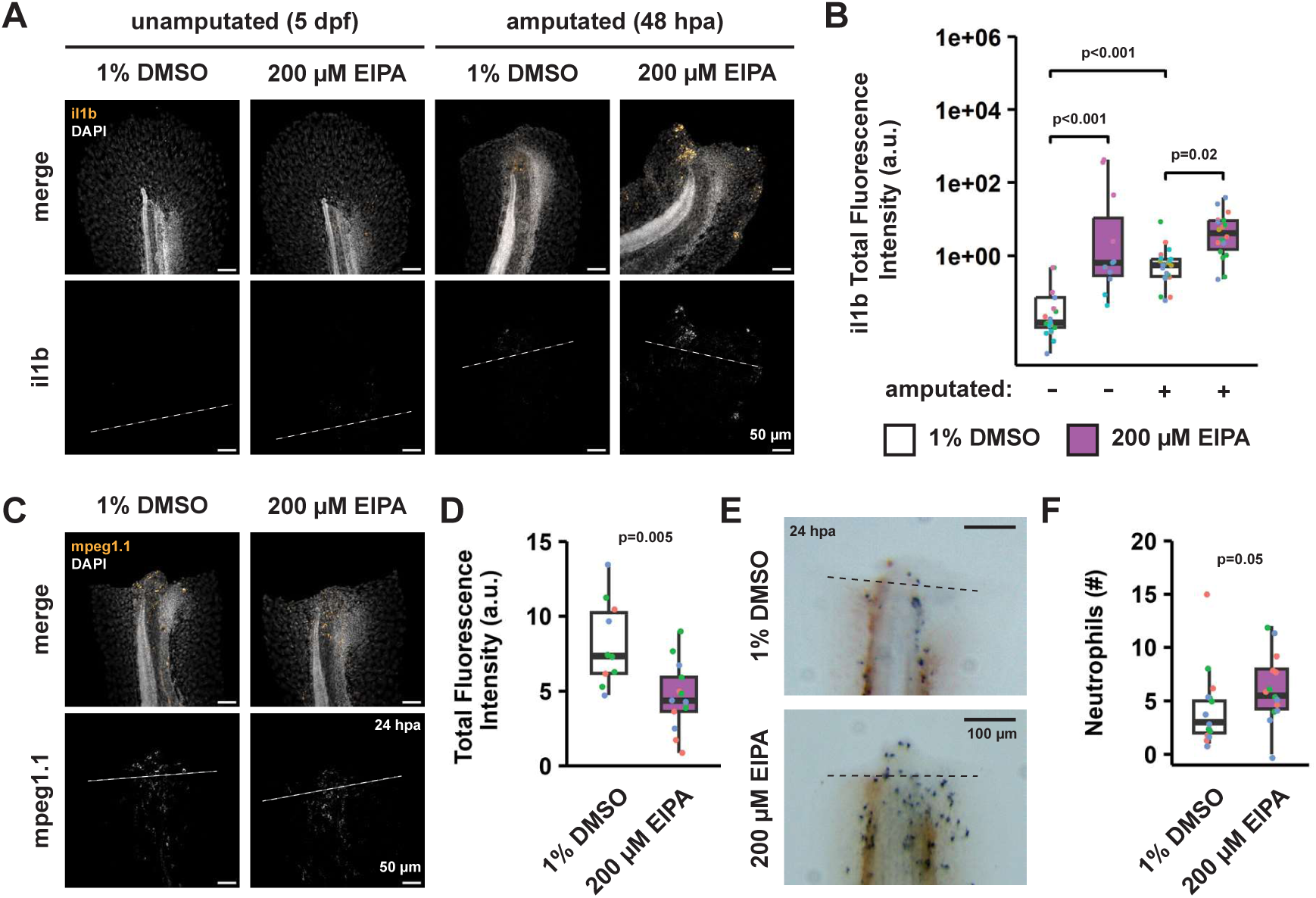
EIPA treatment results in a pro-inflammatory state post-amputation. (A,B) Representative maximum projection images (A) and quantifications (B) of *il1b* RNA-FISH in unamputated and amputated wildtype larval tails. Dotted lines indicate boundaries of ROIs used for quantification. Scale bars: 50 µm. Dots represent fish. Dot colors represent experimental days. Boxplots show median and interquartile range. n=11-22 per group. p-values from Kruskal-Wallis (p<0.001) and Dunn’s tests. (C,D) Representative maximum projection images (C) and quantifications (D) of *mpeg1.1* RNA-FISH in amputated wildtype larval tails at 24 hpa. Dotted lines indicate boundaries of ROIs used for quantification. Dots represent fish. Dot colors represent experimental days. Boxplots show median and interquartile range. n=10,13. p-value from t-test. (E,F) Representative images (E) and quantifications (F) of Sudan black staining for neutrophils in amputated wildtype larval tails at 24 hpa. Dotted lines indicate boundaries of ROIs used for quantification. Dots represent fish. Dot colors represent experimental days. Boxplots show median and interquartile range. n=15,14. p-value from Wilcoxon rank sum test.

We next examined the effects of EIPA treatment on myeloid cells, which migrate toward the wound site upon amputation and regulate the balance between pro- and anti-inflammatory signaling (Hasegawa et al., 2017; Li et al., 2012; Nguyen-Chi et al., 2017). We visualized the macrophage marker *mpeg1.1* (Ellett et al., 2011) by RNA-FISH at 24 hpa and found that EIPA treatment led to decreased levels of *mpeg1.1* puncta near the wound relative to DMSO-treated controls, suggesting a reduced number of macrophages in the wound region (Fig. 5C,D). We also quantified neutrophils at 24 hpa by Sudan black staining and found increased numbers of neutrophils near the wound in EIPA-versus DMSO-treated larvae (Fig. 5E,F). Consistent with this finding, our RNA-seq data at 48 hpa indicated that the neutrophil marker *mpx* (Bennett et al., 2001) was upregulated (log_2_FC 0.64) in amputated larvae following EIPA treatment, further supporting an increased number of neutrophils in the wound region (Fig. 3D). Thus, EIPA treatment alters the balance of myeloid cells in the wound region at 24 hpa, increasing neutrophils and decreasing macrophages. Given the pro-inflammatory role of neutrophils and anti-inflammatory role of macrophages at later stages of tail regeneration, these results suggest that EIPA treatment in amputated larvae promotes inflammation via opposing effects on macrophage and neutrophil populations in the wound region.

We next determined whether EIPA treatment disrupts the behavior of neutrophils, the earliest myeloid cells to arrive at the wound site (Li et al., 2012; Mathias et al., 2006). We used *in vivo* time lapse imaging to track the migration of neutrophils over the first 4 hpa in DMSO- or EIPA-treated *Tg(mpx:dendra2); Tg(cdh1:cdh1-tdTomato)* larvae (Cronan et al., 2018; Yoo and Huttenlocher, 2011) (Fig. 6A). In DMSO-treated larvae, neutrophils were observed migrating toward the amputation plane upon wounding (Fig. 6A; Supp Movies 1, 2). In EIPA-treated larvae, we saw no difference in the number of neutrophils tracks (Fig. 6B; Supp Movies 3, 4) or the “success fraction” of migrating neutrophils that arrived at the wound edge by 4 hpa (Fig. 6C). These data indicate that EIPA treatment does not reduce the initial size of the recruited neutrophil population, nor does it prevent neutrophil directed migration post-amputation.

**Fig. 6.**
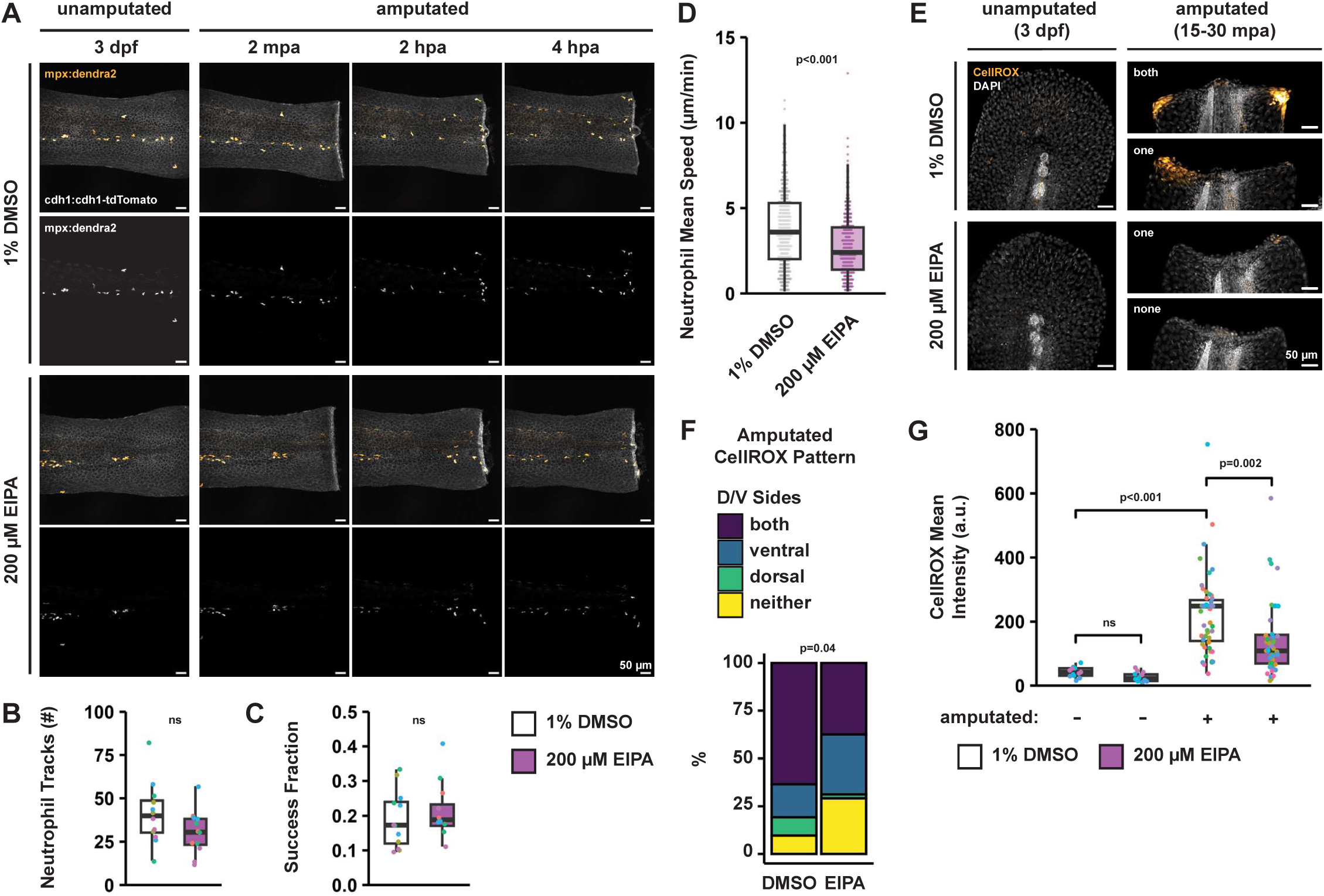
EIPA treatment disrupts neutrophil dynamics and ROS production post-amputation. (A) Representative maximum projection images of *Tg(mpx:dendra2); Tg(cdh1:cdh1-tdTomato)* larval tails from pre-amputation to 4 hpa. Scale bars: 50 µm. (B,C) Number of neutrophil tracks (B) and success fraction (i.e. fraction of migrating neutrophils that arrived at the wound edge by 4 hpa) (C) in amputated *Tg(mpx:dendra2); Tg(cdh1:cdh1-tdTomato)* larval tails. Dots represent fish. Dot colors represent experimental days. Boxplots show median and interquartile range. N=12 fish per group. p-values from t-tests. (D) Neutrophil mean speed over 4 hpa quantified in amputated *Tg(mpx:dendra2); Tg(cdh1:cdh1-tdTomato)* larval tails. Dots represent cells. Boxplots show median and interquartile range. n=494,365 cells. N=12 fish per group. p-value from Wilcoxon rank sum test. (E) Representative maximum projection images of CellROX staining for ROS in unamputated and amputated wildtype larval tails. Scale bars: 50 µm. (F) Distribution of CellROX patterns in amputated wildtype larval tails. n=51,47. p-value from Fisher’s Exact test. G) CellROX mean intensity quantified in unamputated and amputated wildtype larval tails. Dots represent fish. Dot colors represent experimental days. Boxplots show median and interquartile range. n=13,14 (unamputated groups) or n=51,47 (amputated groups). p-values from Kruskal-Wallis (p<0.001) and Dunn’s tests.

Despite these similarities, we found that the mean speed of each neutrophil over time was reduced with EIPA relative to DMSO (Fig. 6D). This finding prompted us to ask whether EIPA treatment might impact neutrophil reverse migration, despite it not having a measurable effect on neutrophil migration toward the wound. Specifically, for the subset of neutrophils that started off close to amputation plane, we tracked the total motion over the time of observation along the anterior/posterior (Fig. S10A) and dorsal/ventral axes (Fig. S10B). The mean end positions at 4 hpa were not altered by EIPA treatment, but the spread of end positions was narrower for all directions in EIPA-treated larvae relative to DMSO-treated controls, suggesting that EIPA treatment leads to a decrease in the net movement of these neutrophils in all directions. These results support the conclusion that EIPA treatment reduces neutrophil speed without affecting directionality. We note that the net slower migration speed may delay resolution of inflammation simply because neutrophils in EIPA-treated larvae take longer to leave the amputation site, perhaps contributing to the excess neutrophil accumulation and increased inflammation that we observed at 24 hpa (Fig. 5E,F).

We next asked whether EIPA treatment disrupts signals that recruit myeloid cells to the wound. One such key signal is ROS, which is produced by epithelial cells at ∼20 mpa and is required for tail regeneration (de Oliveira et al., 2014; Ding et al., 2025; Niethammer et al., 2009; Romero et al., 2018; Sipka et al., 2021; Tauzin et al., 2014; Yoo et al., 2012). We quantified ROS production using the dye CellROX. While the CellROX signal was minimal in unamputated larvae, it was observed in DMSO-treated larvae at 15 to 30 mpa in the fin fold close to the wound site—on both sides of the notochord in 63% of tails, on only the ventral side in 17% of tails, and on only the dorsal side in 10% of tails (Fig. 6E-G; Fig. S10C). We suspect that these different patterns result from differing ROS kinetics in the dorsal and ventral fin folds, with greater or more persisting signal on the ventral side next to the caudal vein. By contrast, in EIPA-treated amputated larvae the distribution of CellROX signal was shifted to 38% of tails with signal on both sides of the notochord, 31% of tails with signal on only the ventral side, 2% of tails with signal on only the dorsal side, and 29% of tails with no signal (Fig. 6E,F; Fig. S10C). EIPA treatment also decreased the CellROX mean intensity overall by 36% (Fig. 6E,G; Fig. S10C). Thus, our data collectively indicate that EIPA treatment results in defects early in the regeneration process, including reduced neutrophil motility and decreased ROS production.

### The GSK3 inhibitor BIO partially rescues tail regeneration in larvae exposed to EIPA or low pHe

Our data indicated that EIPA treatment results in decreased proliferation and increased inflammation during tail regeneration. Increased GSK3 activity is also known to promote decreased proliferation and increased inflammation in many contexts (Souder and Anderson, 2019). Furthermore, inhibiting GSK3 enhances tail regeneration in *Xenopus* tadpoles (Contreras et al., 2009) as well as in both larval (Ouyang et al., 2017) and adult zebrafish (Nachtrab et al., 2011; Sarmah et al., 2019). We therefore tested whether the effects of EIPA on tail regeneration might occur in part through changes in GSK3 activity, with the prediction that EIPA treatment increases GSK3 activity. We tested our prediction by immunolabeling for GSK3 phosphorylated at its inhibitory site Ser21/9 (pGSK3). We quantified the pGSK3 signal in regions adjacent to the notochord bead (an injury-induced extrusion of notochord cells) and found reduced intensity with EIPA compared with DMSO at 48 hpa (Fig. 7A,B). This result suggests that EIPA treatment leads to an aberrant increase in active GSK3 during tail regeneration.

**Fig. 7.**
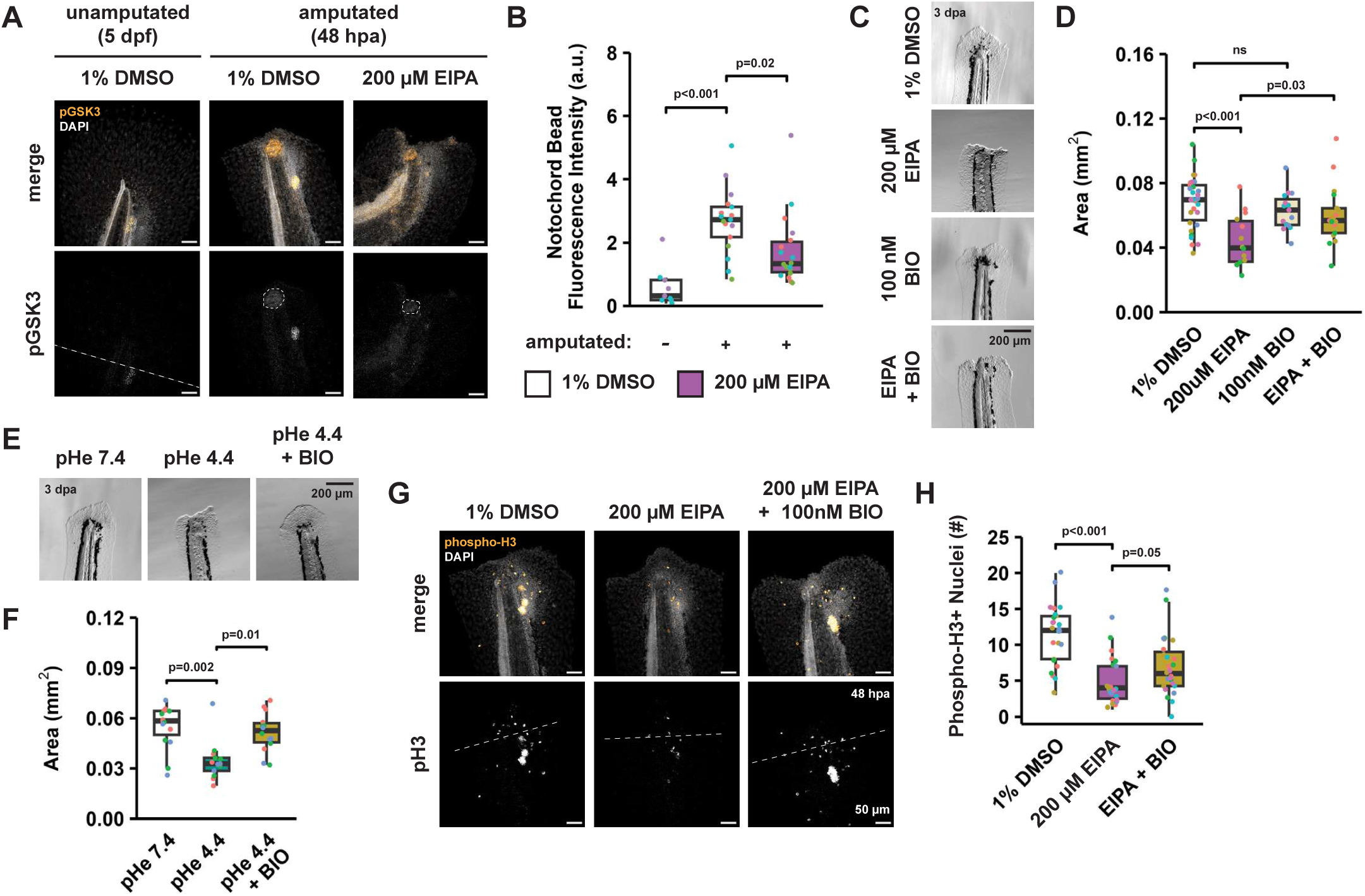
The GSK3 inhibitor BIO partially restores regeneration in larvae exposed to EIPA or low pHe. (A,B) Representative images (A) and quantification (B) of pGSK3 immunolabeling in unamputated and amputated wildtype larval tails. Dotted lines indicate boundaries of ROIs used for quantification. n=9-18 per group. p-values from Kruskal-Wallis (p<0.001) and Dunn’s tests. (C,D) Representative images (C) and area of regrowth (D) of wildtype larval tails amputated in indicated drugs at 3 dpa. n=14-30 per group. p-values from one-way ANOVA (p<0.001) and Tukey HSD tests. (E,F) Representative images (E) and area of regrowth (F) of wildtype larval tails amputated in indicated pHe and drugs at 3 dpa. n=13-15 per group. p-values from Kruskal-Wallis (p<0.001) and Dunn’s tests. (C,E) Scale bars: 200 µm. (G,H) Representative maximum projection images (G) and quantifications (H) of phospho-H3 immunolabeling in amputated wildtype larval tails at 48 hpa. Dotted lines indicate boundaries of ROIs used for quantification. n=23-26 per group. p-values from Kruskal-Wallis (p<0.001) and Dunn’s tests. (B,D,F,H) Dots represent fish. Dot colors represent experimental days. Boxplots show median and interquartile range.

Based on these changes in pGSK3, we tested whether inhibiting GSK3 using the pharmacological agent BIO might rescue regeneration in EIPA-treated larvae. We found that while 100 nM BIO alone had no effect on area of regrowth at 3 dpa, co-treatment with BIO and EIPA partially restored area of regrowth compared with EIPA alone (Fig. 7C,D). BIO treatment also rescued the decrease in area of regrowth caused by exposure to low pHe (Fig. 7E,F). Moreover, BIO rescued the proliferation defect caused by EIPA at 48 hpa, as indicated by phospho-H3 immunolabeling (Fig. 7G,H). Together, these data support a growing body of evidence that GSK3 inhibition is pro-regenerative and suggest that limiting the increase in pHi post-amputation with EIPA or low pHe may impair tail regeneration in part through GSK3-mediated signaling.

## DISCUSSION

We quantified pHi dynamics during zebrafish larval tail regeneration using a newly generated transgenic pHi biosensor line *Tg(ef1α:mKate2-SEpHluorin)* along with calibration methods. Compared with the use of pHi-sensitive dyes (Mölich and Heisler, 2005; Raja et al., 2020; Yew et al., 2020) or injection of mRNA encoding a pHi biosensor (Ermakova et al., 2018; Tarbashevich et al., 2015; Vandestadt et al., 2021), our pHi biosensor line has the advantage of ubiquitous and stable mKate2-SEpHluorin expression, allowing for measurements over longer periods of time. With these advantages, we identified widespread changes in pHi that occur upon zebrafish larval tail amputation, including an initial decrease consistent with a previous report in *Xenopus laevis* (Matlashov et al., 2015), followed by an increase above pre-amputation values that is partially dependent on NHE activity. We found that pHi dynamics is critical during both the wound healing and proliferative phases of zebrafish larval tail regeneration. Moreover, our data suggest distinct functions of pHi dynamics during each phase, including effects on proliferation, inflammation, and GSK3 activity.

We first showed that attenuating the post-amputation increase in pHi impairs cell proliferation at 48 hpa. This could be mediated in part by increased GSK3 activity, consistent with previous evidence that GSK3 inhibits cell cycle progression by directly phosphorylating and destabilizing targets such as cyclin E, cyclin D1, c-myc, and cdc25A (Xu et al., 2009). Decreased ROS levels might also contribute to this effect, as ROS production is required for proliferation during tail regeneration (Yoo et al., 2012). Finally, disrupting pHi might also directly impair cell cycle progression, as shown in cell lines and mouse xenograft models (Flinck et al., 2018; Putney and Barber, 2003; Spear and White, 2023). Our *in vivo* data thus support the conclusion from previous work in clonal cells that increased pHi enables cell proliferation.

We also found that EIPA treatment leads to increased inflammation at 48 hpa, consistent with previous reports that decreasing pHe or pHi in cell lines promotes a pro-inflammatory phenotype (Chae et al., 2023; Kawakami et al., 2024; Li et al., 2023; Rajamäki et al., 2013; Riemann et al., 2016, 2015; Wu et al., 2025). We predict that EIPA treatment promotes inflammation in part through its opposing effects on myeloid populations in the wound region at 24 hpa, increasing the number of neutrophils, which are pro-inflammatory (Li et al., 2012), and decreasing the number of macrophages, which are primarily anti-inflammatory during the proliferative phase (Hasegawa et al., 2017; Nguyen-Chi et al., 2017). We found that EIPA treatment leads to reduced neutrophil motility during the first 4 hpa, which could in part be due to decreased ROS levels, as ROS is an established myeloid chemoattractant (de Oliveira et al., 2014; Niethammer et al., 2009; Tauzin et al., 2014). This result could also be explained by a cell-autonomous effect in neutrophils, as NHE1 activity is required for chemotaxis in human primary neutrophils (Nagy et al., 2024; Oster et al., 2022). An additional possibility is that neutrophils are slowed due to increased GSK3 activity (as described below), which inhibits F-actin remodeling and impairs chemotaxis in mouse primary neutrophils (Tang et al., 2011).

Indeed, we demonstrated that EIPA treatment leads to increased GSK3 activity at 48 hpa, consistent with previous reports in cell lines (Darmellah et al., 2009; Javadov et al., 2009). Furthermore, the GSK3 inhibitor BIO partially rescues regeneration with EIPA or low pHe, similar to previous reports in zebrafish (Nachtrab et al., 2011; Ouyang et al., 2017; Sarmah et al., 2019) and *Xenopus* (Contreras et al., 2009) indicating that GSK3 inhibitors promote tail regeneration. Inhibiting GSK3 activity also promotes regeneration in multiple other systems, including spinal cord (Dong et al., 2023), kidney (Jamadar and Rao, 2020), liver (Bhushan et al., 2017), and muscle (Pansters et al., 2015). Thus, our results support the growing body of evidence that GSK3 inhibition is a pro-regenerative signaling mechanism, further linking it to pHi.

In summary, we identified a previously unreported role for pHi dynamics in zebrafish larval tail regeneration. It remains to be determined whether changes in pHi are more widely implicated in regeneration across species and tissues beyond the tail regeneration model studied here. Similar changes in pHi observed in *Xenopus* tails post-amputation (Matlashov et al., 2015) suggest that pHi dynamics might also influence appendage regeneration in other species. Furthermore, according to one report, pHi decreases upon zebrafish larval spinal cord transection, and incubation in low pHe enhances precursor neuron recruitment to the injury site (Vandestadt et al., 2021). These findings raise the possibility that changes in pHi might play a broad role during regeneration of additional tissues.

Finally, there is evidence for changes in pHi and spatiotemporal regulation of NHE1 expression in human wounds. Studies using a luminescent pHe sensor on human skin reported patterns in pHe and NHE1 expression around the center of wounds (Schreml et al., 2014, 2011). These features are also dynamic, with pHe decreasing and NHE1 expression increasing over the course of wound healing (Hachem et al., 2005; Haverkampf et al., 2017). In conclusion, our findings establish a previously unrecognized functional role for pHi dynamics during zebrafish larval tail regeneration, with significance to the regeneration field as well as relevance to human regenerative medicine.

## MATERIALS AND METHODS

### Zebrafish Husbandry

For experiments conducted in the Barber and Wagner labs, all zebrafish (wildtype AB background) were handled according to Protocol AN199988 approved by the University of California, San Francisco Institutional Animal Care and Use Committee. Embryos and larvae were incubated in 0.3X Danieau medium (17.4 mM NaCl, 0.21 mM KCl, 0.12 mM MgSO_4_, 0.18 mM Ca(NO_3_)_2_, 1.5 mM HEPES, pH 7.6).

For neutrophil tracking experiments conducted in the Theriot lab, all zebrafish (wildtype TAB5 background) were handled according to Protocol 4427-01 approved by the University of Washington Institutional Animal Care and Use Committee. Embryos and larvae were incubated in E3 medium (5 mM NaCl, 0.17 mM KCl, 0.33 mM CaCl_2_, 0.33 mM MgSO_4_).

All adult zebrafish were housed according to standard procedures on a 14 h light, 10 h dark cycle at 28.5°C. Crosses were set up by natural spawning. All experiments were conducted on 3 to 6 dpf larvae, at which stage sex determination is not yet possible. All embryos and larvae were housed in Petri dishes in incubators maintained at 28.5°C.

### Zebrafish Transgenic Lines

To generate the *Tg(ef1α:mKate2-SEpHluorin)* transgenic line, an mKate2-SEpHluorin cassette generated in the Barber lab (mKate2 sequence from Addgene #31869 and SEpHluorin sequence from Addgene #32001) was cloned by Gibson Assembly (New England Biolabs, E2611) into a pMTB vector backbone (Wagner et al., 2018) containing Tol2 elements for genomic integration. Wildtype AB embryos were injected at the 1-cell stage with 1 to 2 nL of injection solution containing 10 ng/µL plasmid, 50 ng/µL Tol2 integrase mRNA, and 0.05% phenol red (Sigma, P0290).

The *Tg(mpx:dendra2)* and *Tg(cdh1:cdh1-tdTomato)* lines used for neutrophil tracking were obtained from the Huttenlocher lab (Yoo and Huttenlocher, 2011) and Cronan lab (Cronan et al., 2018), respectively. Both lines were maintained in a wildtype TAB5 background.

### pHi Measurements & Calibrations

For pHi measurements, *Tg(ef1α:mKate2-SEpHluorin)* larvae were imaged live on a Yokogawa CSU-W1/SoRa spinning disk confocal microscope equipped with an ORCA Fusion BT sCMOS camera (Hamamatsu). 3 dpf larvae were pre-incubated for 1 h in indicated treatment solution (1% DMSO vehicle control, 200 µM EIPA, pHe 7.4, or pHe 4.4), then transferred to indicated treatment solution plus 0.1 mg/mL Tricaine (Sigma, A5040). For pre-amputation imaging, larvae were mounted on a 35 mm MatTek dish with 20 mm glass microwell (MatTek P35G-1.5-20-C) with rectangular agarose wells. For post-amputation imaging, larvae were removed from the microscope, amputated, and mounted in 1% low melting point agarose (Sigma, A0701) on a MatTek dish for imaging. Images were acquired as Z-stacks of 2 µM optical sections using an Apo LWD Lambda S 40X objective (Nikon). Larvae were maintained at 28.5°C throughout imaging.

For calibrating SEpHluorin/mKate2 ratios to pHi values, 3 dpf amputated *Tg(ef1α:mKate2-SEpHluorin)* larvae were incubated for 20 min in calibration buffer (25 mM HEPES, 105 mM KCl, 1 mM MgCl_2_, 40 µM nigericin (Invitrogen, 1495), 0.1 mg/mL Tricaine, and 1% DMSO, with pH values set to range from 6.8 to 7.8 using HCl and KOH). Larvae were imaged in a MatTek dish with agarose wells as described above.

Images were processed using NIS-Elements software. Background was subtracted using an empty ROI from each optical section, and pseudo-colored ratiometric images were generated using the Ratio View function. Ratios were quantified using Imaris software. A surface was created using the mKate2 channel and cut into 40 µM slices from the wound site. Ratios were calculated using the Channel Arithmetics function.

### Regeneration Assays

3 dpf larvae were anesthetized with 0.1 mg/mL Tricaine and placed on a clean 100 mm Petri dish lid where they were amputated with a clean razor blade. Tail amputations were performed at the posterior end of the pigment gap (Romero et al., 2018). Amputated larvae were then immediately transferred to autoclaved 60 x 15 mm glass dishes (Corning, 3160-60) for recovery (max 8 larvae per dish). At 3 dpa (6 dpf), larvae were imaged on a Leica M80 stereomicroscope. Area of tissue regrowth was measured using FIJI software (Schindelin et al., 2012), with images blinded by the Blind Analysis Tools plugin. Area was measured based on an ROI drawn around the presumptive regenerated tissue starting at the anterior boundary of the notochord bead through the end of the fin fold (amputated tails) or around tail tissue starting at the posterior end of the pigment gap through the end of the fin fold (unamputated tails).

### Drug Treatments

The following drugs were used: 5-(N-Ethyl-N-isopropyl)amiloride (EIPA) (Enzo, ALX-550-266-M005, Lots 44GV48 and 45FV08) stored at -20°C as 50 mM stocks in DMSO (EIPA dose-response and survival curves should be determined for each vendor and lot because we observed discrepancies when testing EIPA from different vendors); atorvastatin (Cayman, 10493) stored at -20°C as 25 mM stocks in DMSO; (2’Z,3’E)-6-Bromoindirubin-3′-oxime (BIO) (Sigma, B1686) stored at -80°C as 10 mM stocks in DMSO. For experiments, drug dilutions were prepared fresh and supplemented with DMSO as needed to bring all solutions to a final DMSO concentration of 1%.

### pHe Manipulation

Media of different pHe were made by adding HCl or NaOH to 0.3X Danieau supplemented with 20 mM HEPES. NaCl was added as needed to make all media used in a given experiment isosmotic. Final osmolarity was ∼100 mOsm/L. Medium was changed daily throughout experiments.

### CRISPR/Cas9 Genome Editing

CRISPR-Cas9 genome editing was conducted according to the Integrated DNA Technologies (IDT) Alt-R CRISPR-Cas9 protocol. Guide RNAs were generated by annealing 100 µM universal tracrRNA (IDT) and 100 µM custom crRNA (Table 1) diluted fresh 1:15 in nuclease-free duplex buffer (IDT) for 5 min at 95°C. Ribonucleoprotein (RNP) complexes were then assembled by mixing equal volumes of gRNA and Cas9 protein (IDT, 1081060) diluted fresh to 1.5 µg/µL in Cas9 working buffer (20 mM HEPES, 150 mM KCl, pH 7.5) and incubating for 10 min at 37°C. Wildtype AB embryos were injected at the 1-cell stage with 1 to 2 nL of injection solution consisting of freshly prepared RNP complex and 0.05% phenol red (Sigma, P0290). Injected embryos were used for regeneration assays in the F_0_ crispant screen. Editing was confirmed in amputated larvae by polymerase chain reaction (PCR) (30 sec extension time; primers listed in Table 2) followed by the T7 endonuclease I (T7E1) mismatch cleavage assay (Alt-R Genome Editing Detection Kit, IDT, 1075932).

To generate stable knockout lines, injected embryos were raised to adulthood and outcrossed to wildtype AB embryos. F_1_ fish were incrossed in pairs, and their F_2_ progeny were screened by T7E1 assay and Sanger sequencing to identify edited alleles with a premature stop codon. Regeneration assays were conducted by incrossing F_1_ parents confirmed to be heterozygous for the same mutant allele, and amputated larvae were genotyped by T7E1 assay or Sanger sequencing.

### Protein Sequence Alignment

Protein sequence alignments were performed using the Uniprot Align feature (Ahmad et al., 2025) using the following Entry IDs: P19634, A0A8M2BKS9, A9XP99, Q9UBY0, A0AB32U1D4, Q9Y2E8, Q5PR36.

### Bulk RNA Sequencing

Bulk RNA-sequencing was conducted on three biological replicates per condition, each consisting of pooled tail tissue from 30 wildtype AB larvae treated and amputated on the same days. Tail tissue was isolated by dissection on ice, with amputated tails cut approximately 200 to 250 µM from the wound site and unamputated tails cut at the anterior end of the pigment gap. Tail chunks were placed individually in PCR strip tube caps, and caps were inverted into PCR strip tubes containing 50 µL TRIzol (Invitrogen, 15596026) each. Tissue was partially dissociated by pipetting before TRIzol samples were pooled and further dissociated with a 30G syringe. Samples were vortexed and stored at -80°C.

Total RNA was extracted from TRIzol samples using 0.5 volumes chloroform. RNA was precipitated from the resulting aqueous layer overnight at -80°C using 1 volume ice cold isopropanol, 0.1 volumes ice cold 3M sodium acetate (Invitrogen AM9740), and 0.01 volumes Glycoblue (Invitrogen, AM9515). RNA samples were washed 3X with 70% ice cold ethanol and resuspended in 15 µL nuclease-free water. Finally, RNA samples were submitted to Novogene for QC, library preparation by poly-A capture, and 150 bp paired-end sequencing (Illumina X Plus).

Data were processed using the nf-core/rna-seq pipeline (v3.12.0) (Ewels et al., 2020). Pseudo-alignment by Salmon (Patro et al., 2017) was specified, using the GRCz11/danRer11 zebrafish reference genome and the Lawson lab transcriptome annotation (v4.3.2) (Lawson et al., 2020). Default parameters were otherwise used. Raw and processed data are available through the Gene Expression Omnibus repository (GSE318689).

Downstream analysis was conducted using R (v4.3.3). Differential gene expression analysis was performed using DESeq2 (v1.42.1) (Love et al., 2014), with differentially expressed genes defined by adjusted p-value < 0.05 at 6 hpa and by both adjusted p-value < 0.05 and |log_2_ fold change| > 0.5 at 48 hpa. Gene ontology analysis was performed using the enrichGO function from clusterProfiler (v4.10.1) (Wu et al., 2021).

### Fixation & Permeabilization

For immunolabeling, EdU, TUNEL, and RNA-FISH, larvae were fixed in 4% paraformaldehyde (PFA) in PBS overnight at 4°C, washed 3X in PBS supplemented with 0.1% Tween-20 (PBST), series dehydrated into methanol, and stored minimum overnight at -20°C. On Day 1 of staining, samples were series rehydrated into 0.1% PBST, and tail tissue was isolated by dissection. Tail tissue was permeabilized in 7.5 µg/µL Proteinase K (Sigma, P556) for 15 min, washed 2X in 0.1% PBST, refixed in 4% PFA for 15 min at RT, and washed 3X in either PBS supplemented with 1% Triton X-100 (PBTx) or 0.1% PBST.

### Immunolabeling

Samples were incubated on Day 1 in blocking solution (5% normal goat serum (Thermo Fisher, 10000C) in 1% PBTx) for 1 to 2 h. Samples were then incubated in primary antibody diluted 1:300 in blocking solution at 4°C (rabbit anti-phospho-H3 (Ser10) (Sigma, 06-570) for one overnight or rabbit anti-pGSK3α/β (Ser21/9) (Cell Signaling, 9331) for three overnights). On Day 2, samples were washed 8X (15 min/wash) in 1% PBTx and incubated in secondary antibody diluted 1:500 in blocking solution overnight at 4°C (goat anti-rabbit IgG Alexa Fluor 647 (Invitrogen, A-21245) at 1:500). On Day 3, samples were washed 6X (10 min/wash) in DAPI (Invitrogen, D1490) diluted 1:1000 in 1% PBTx for 1 to 2 h and mounted on slides with ProLong Gold Antifade Mountant (Invitrogen, P36930) for imaging as described below.

### EdU Staining

EdU staining was conducted using the Click-iT EdU Cell Proliferation Kit, Alexa Fluor 488 (Invitrogen, C10337). Amputated larvae were soaked at ∼45 hpa in 400 µM EdU in a 12-well plate for 45 min at 28.5°C. Larvae were fixed and permeabilized as described above. Fixed samples were incubated in Click-iT reaction cocktail for 30 min at RT. Samples were then washed 6X (10 min/wash) in DAPI diluted 1:1000 in 1% PBTx for 1 to 2 h, followed by immunolabeling or mounting on slides with ProLong Gold Antifade Mountant for imaging as described below.

### TUNEL Labeling

Samples were washed in PBS and incubated in TUNEL Enzyme Solution diluted 1:100 in TUNEL Label Solution (*In Situ* Cell Death Detection Kit, TMR red, Roche, 12156792910) for 1 h at 37°C. Special care was taken to always maintain samples submerged in liquid. Samples were then washed 3X in 0.1% PBST, incubated in DAPI diluted 1:1000 in 0.1% PBST for 1 to 2 h, and mounted on slides with ProLong Gold Antifade Mountant for imaging as described below.

### RNA Fluorescence *In Situ* Hybridization (RNA-FISH)

RNA-FISH was conducted according to the Molecular Instruments protocol for whole-mount zebrafish larvae using hybridization chain reaction v3.0 probes and kits. Probes for *dlx5a*, *aldh1a2*, *il1b*, and *mpeg1.1* were custom ordered from Molecular Instruments. Briefly, on Day 1, samples were pre-hybridized in probe hybridization buffer for 30 min at 37°C and incubated in probe solution (5 pmol of each probe in 250 µL probe hybridization buffer) overnight at 37°C. On Day 2, samples were washed in probe wash buffer, washed in 5X SSCT, pre-amplified in amplification buffer for 30 min at RT, and incubated in hairpin solution (15 pmol of each snap-cooled hairpin in 250 µL amplification buffer) overnight at RT. On Day 3, samples were washed in 5X SSCT, washed in DAPI diluted 1:1000 in 5X SSCT for 1 to 2 h, and mounted on slides with ProLong Gold Antifade Mountant for imaging as described below.

### Reactive Oxygen Species Measurements

ROS were measured using CellROX Green Reagent dye (Invitrogen, C10444). 3 dpf wildtype AB larvae were pre-incubated for 1 h in 10 µM CellROX with 1% DMSO or 200 µM EIPA prior to tail amputation. For each condition, 3 to 5 larvae were amputated (∼15 min), incubated in fresh CellROX solution with DMSO or EIPA in a 12-well plate for 15 min at 28.5°C, and fixed in 4% PFA in PBS for 1 h at RT. Samples were washed 3X in 0.1% PBST, and tail tissue was isolated by dissection. Tail tissue was washed in DAPI diluted 1:1000 in 0.1% PBST for a 1 to 2 h and mounted on slides with ProLong Gold Antifade Mountant for imaging as described below.

### Fixed Tissue Imaging & Analysis

For immunolabeling, EdU, TUNEL, RNA-FISH, and ROS experiments, images were acquired on a Leica Stellaris 5 laser scanning confocal microscope as Z-stacks of 2 µM optical sections using an HC FLUOTAR L 25x/0.95 W VISIR objective (Leica).

Immunolabeling, EdU, TUNEL, *aldh1a2*, *il1b*, and *mpeg1.1* staining were analyzed using Imaris software. phospho-H3, EdU, and TUNEL signal were quantified by counting spots. *aldha2*, *il1b*, *mpeg1.1*, and pGSK3 signal were quantified by creating a surface and measuring the total surface fluorescence intensity. ROIs were designated using the DAPI channel. For pGSK3 quantification, an ROI was drawn around the notochord bead. For all other quantifications in amputated tails, an ROI was selected starting at the anterior boundary of the notochord bead through the end of the fin fold. For all quantifications in unamputated tails, an ROI was selected to include 100 µm of notochord through the end of the fin fold.

*dlx5a* and ROS staining were analyzed using FIJI software. Images were blinded using the Blind Analysis Tools plugin and analyzed as sum intensity projections. *dlx5a* raw integrated density was measured within an ROI drawn around the presumptive wound epithelium using the DAPI channel (amputated tails) or a 20 µM wide rectangular ROI placed at the posterior end of the pigment gap (unamputated tails). ROS mean intensity was measured within a 100 µM wide rectangular ROI placed at the wound edge (amputated tails) or at the posterior end of the pigment gap (unamputated tails).

### Neutrophil Tracking

Larvae were obtained for neutrophil tracking by crossing the *Tg(mpx:dendra2)* and *Tg(cdh1:cdh1-tdTomato)* lines. Larvae were screened for both transgenes at 3 dpf and prepared for imaging as previously described (Kennard et al., 2021). Briefly, larvae were anesthetized in E3 + Tricaine (E3 + 160 mg/L Tricaine (Sigma, E10521) + 1.6 mM Tris pH 7) and mounted in 1.5% agarose (Invitrogen) in 35 mm #1.5 glass-bottom dishes (CellVis D35-20-1.5N and D35C4-20-1.5N). Larvae were mounted with the *xy* (sagittal) plane parallel to the coverslip, and excess agarose was trimmed from around the tail. Live imaging was performed at RT, variable but measured at 21 to 23°C.

Images were acquired on a Nikon Ti2 inverted microscope with a Piezo Z stage (Applied Scientific Instruments PZ-2300-XY-FT) attached to a Yokogawa CSU-W1 spinning-disk confocal with Borealis attachment (Andor). Illumination was supplied by a laser launch (Vortran VersaLase) with 10 mW 488 nm and 5 mW 561 nm diode lasers (Vortran Stradus). Filters used included a 405/488/561/640/755 penta-band dichroic (Andor), a 488/561 dual-band emission filter (Chroma ZET488/561m) for rapid dual-color imaging, and single-band GFP, RFP, and far-red filters for single-color imaging (Chroma 535/50m, 595/50m, and 700/75 respectively). Images were acquired as Z-stacks of 2-3 µM optical sections using an Apo 20x NA 0.8 air objective (Nikon) and a back-thinned EMCCD camera (Andor iXon EM+) in 16-bit mode without binning. Equipment was controlled using MicroManager (v2.0.1). Larvae were imaged pre-amputation every 2 min for 20 min and post-amputation every 2 min for 4 h.

Images were processed and analyzed using FIJI software (v2.26.0/1.54p). Timelapse images were made into maximum intensity projections and registered using the HyperStackReg 5.6 plugin (Sharma, n.d.). Cell tracking was performed using the Trackmate plugin (Ershov et al., 2022; Tinevez et al., 2017). Cell detection uniformly employed a Laplacian of Gaussian method, with the estimated blob radius of 15 µm for neutrophils. No feature penalty was applied, and track merging and splitting was disallowed. The linking radius for consecutive spots was uniformly set at 75.0 µm. A maximum jump distance of 75.0 µm was permitted, and tracks could bridge up to 4 missing frames. Tracks shorter than a 30 min duration were excluded from analysis. The subset of neutrophils that started off close to the amputation plane was determined by visually examining the track densities and defining a wound region for each fish.

### Sudan Black Staining

Sudan Black protocol was adapted from Levraud et al., (2008). Amputated larvae were fixed at 24 hpa in 4% PFA in PBS for 1 h at RT and washed 3X in 0.1% PBST. Samples were stained in Sudan Black working solution (0.036% Sudan Black (Sigma, 199664) in 70% ethanol) for 1 to 5 h at RT, until staining was visible under a dissection microscope. Samples were washed 3X in 70% ethanol and 1X in 30% ethanol in 0.1% PBST. Samples were cleared in freshly prepared 1% H2O2, 1% KOH for 10 min at RT, then washed 2X in 0.1% PBST. Samples were imaged on a Leica M80 stereomicroscope. Sudan spots were counted manually using FIJI software to blind images (Blind Analysis Tools plugin).

### Statistics

Statistical analysis was conducted using R (v4.3.3). Groups were tested for normality (Shapiro-Wilk test) and homoscedasticity (Bartlett’s test), and tests for the equality of means were chosen accordingly (two-way ANOVA, one-way ANOVA, or Kruskal-Wallis with post-hoc tests).

## ACKNOWLEDGEMENTS

We thank Dr. Torsten Wittmann, the UCSF Biological Imaging Development Colab, and their NIH shared equipment grant S10OD028611-01 for invaluable access to their SoRa spinning disk confocal microscope. We thank Christopher P. Chen for help using the nf-core/rna-seq pipeline to process bulk RNA-seq data. This work was supported by the following: NIH F31 DE032574 award to C.C-F.; Howard Hughes Medical Institute (HHMI) support to J.A.T.; NIH DP2 GM146258, Kinship Foundation Searle Scholar, and NIH R00 GM121852 awards to D.E.W.; and NIH R01 CA197855 award to D.L.B. This article is subject to HHMI’s Immediate Access to Research policy. HHMI lab heads have previously granted a nonexclusive CC BY 4.0 license to the public and a sublicensable license to HHMI in their research articles. Pursuant to those licenses, the author-accepted manuscript of this article can be made freely available under a CC BY 4.0 license immediately upon publication.

