## Supplemental Figures for "Intracellular pH dynamics promotes zebrafish larval tail regeneration"

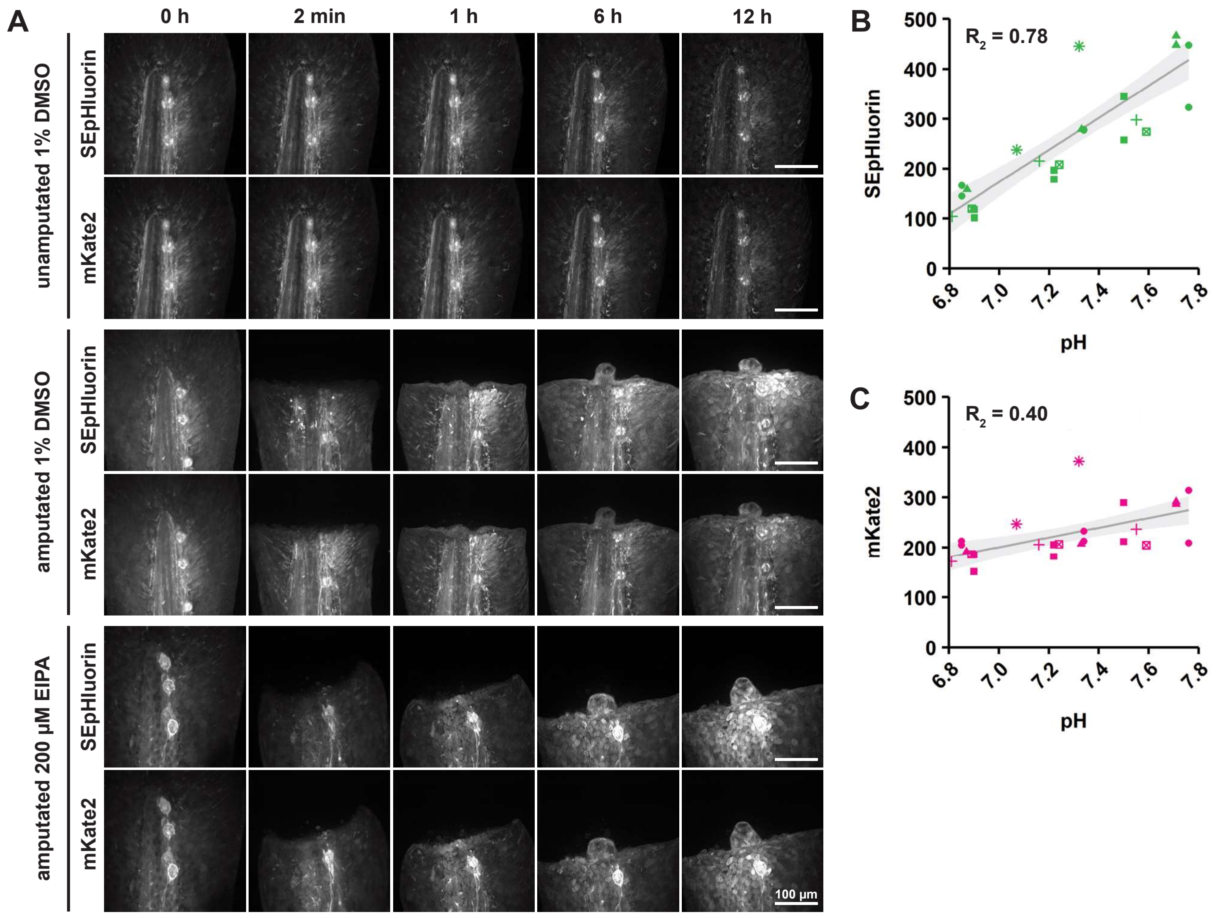


**Fig. S1 - mKate2 and SEpHluorin dynamics**

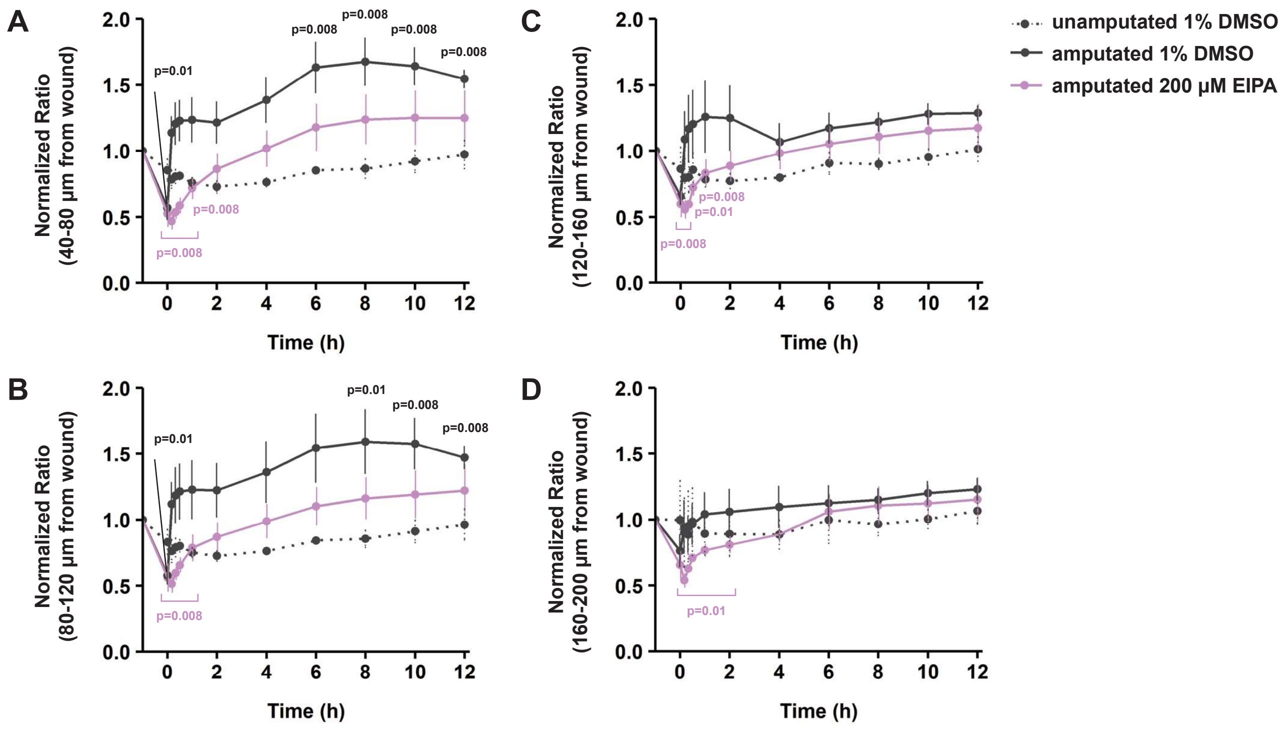


**Fig. S2 - pHi dynamics post-amputation persists up to 120 μm from the wound site**

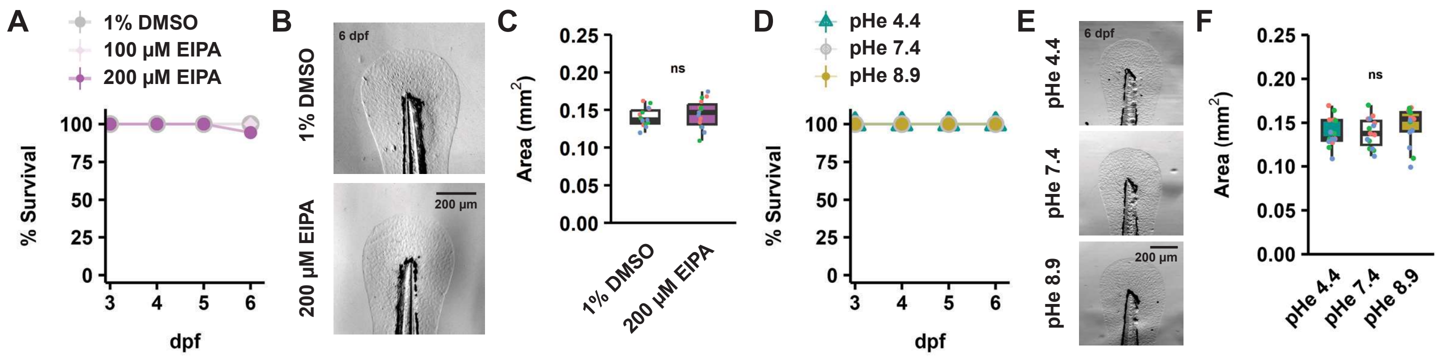


**Fig. S3 - Exposure to EIPA or low pHe has no effect on larval survival or tail growth**

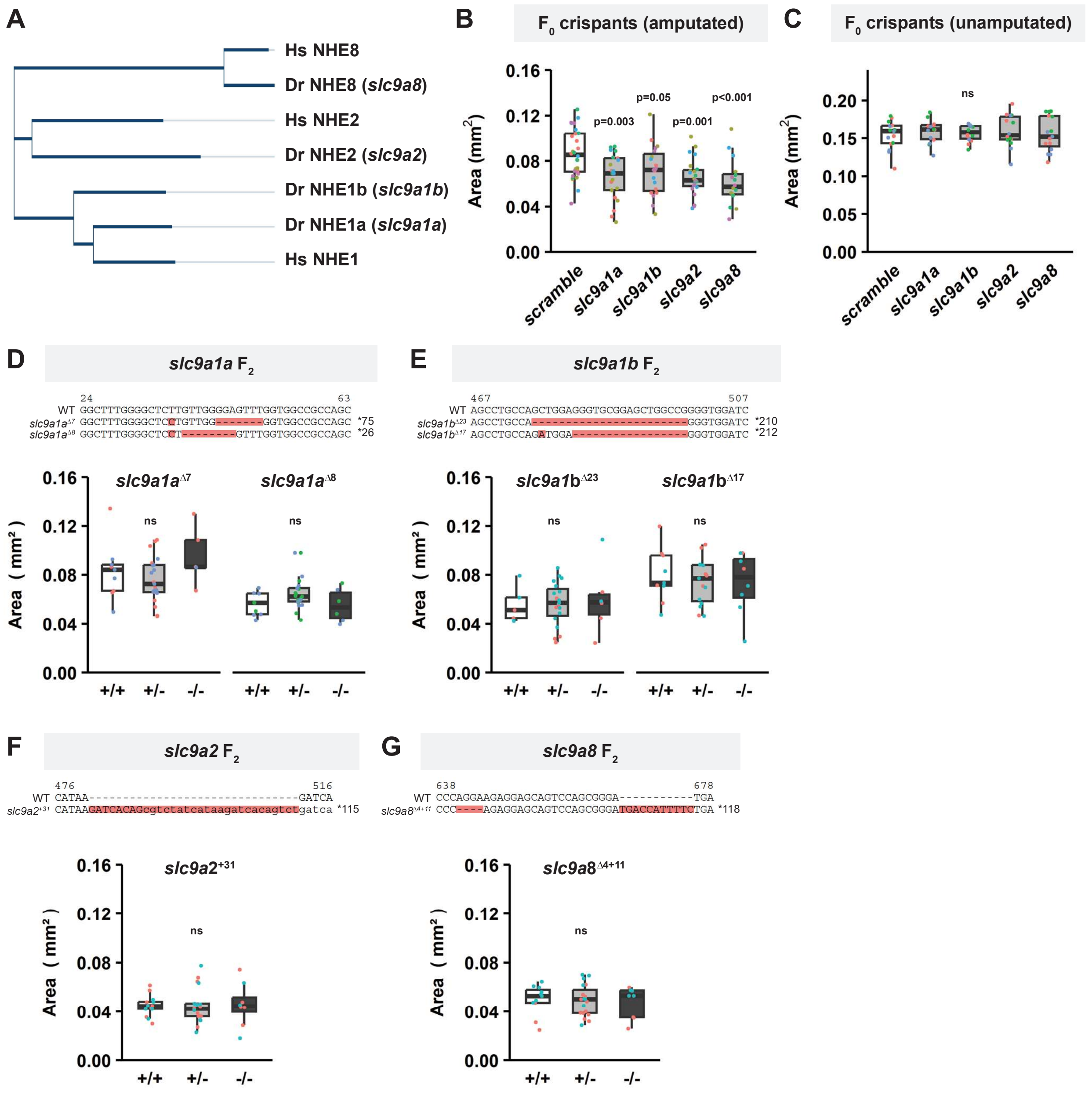


**Fig. S4 - CRISPR/Cas9-mediated knockout of NHE family member genes**

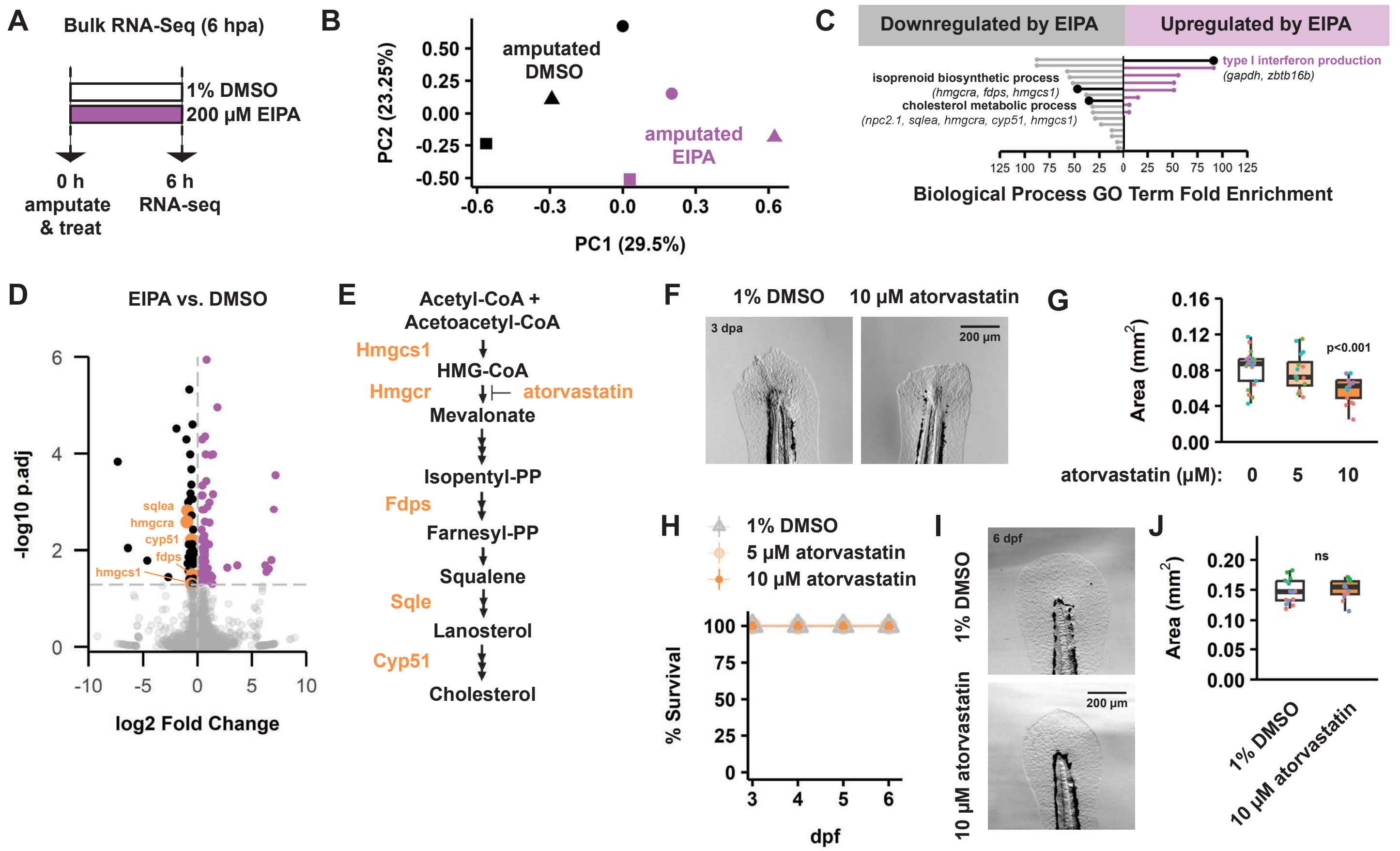


**Fig. S5 - Bulk RNA-seq at 6 hpa identifies role for cholesterol biosynthesis during tail regeneration**

**A**

| Comparison | Interpretation | DEGs | |
|---|---|---|---|
| | | Down | Up |
| Amputated: EIPA vs. DMSO | Effect of EIPA in amputated larvae | 750 | 1421 |
| Amputated: EIPA washout vs. DMSO | Effect of EIPA washout in amputated larvae | 24 | 27 |
| Unamputated: EIPA vs. DMSO | Effect of EIPA in unamputated larvae | 1682 | 2343 |
| DMSO: amputated vs. unamputated | Regeneration in DMSO | 4269 | 4135 |
| EIPA: amputated vs. unamputated | Regeneration in EIPA | 3361 | 3156 |

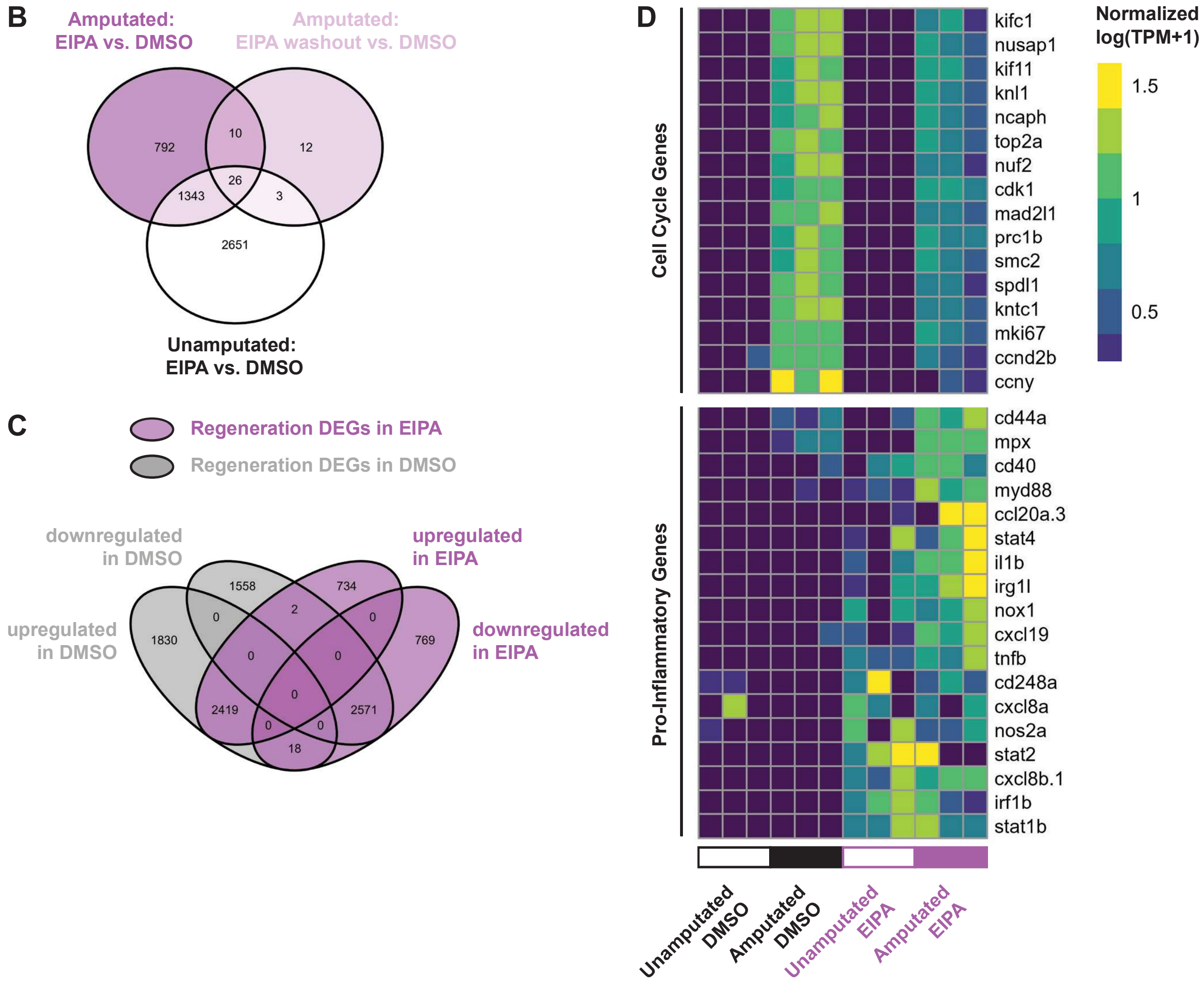


**Fig S6 - Summary of bulk RNA-seq at 48 hpa**

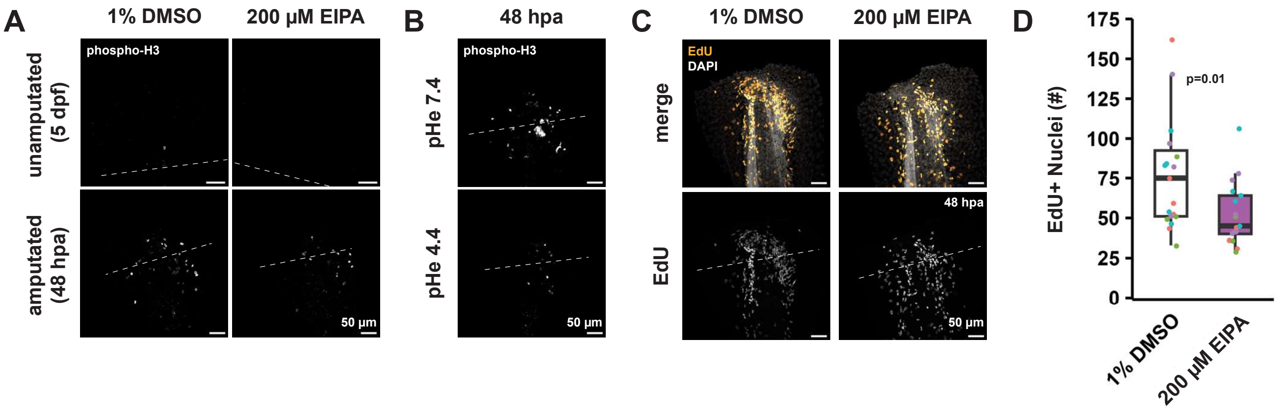


**Fig S7 - Single-channel phospho-H3 images and EdU labeling with EIPA**

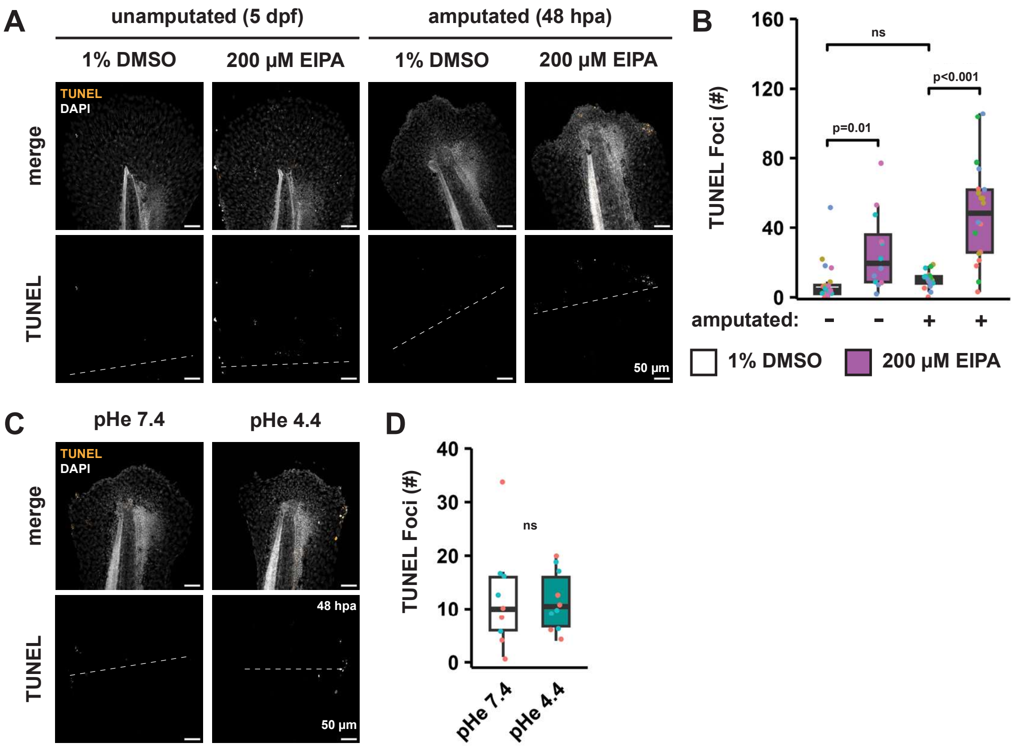


**Fig S8 - Exposure to EIPA but not low pHe leads to increased cell death**

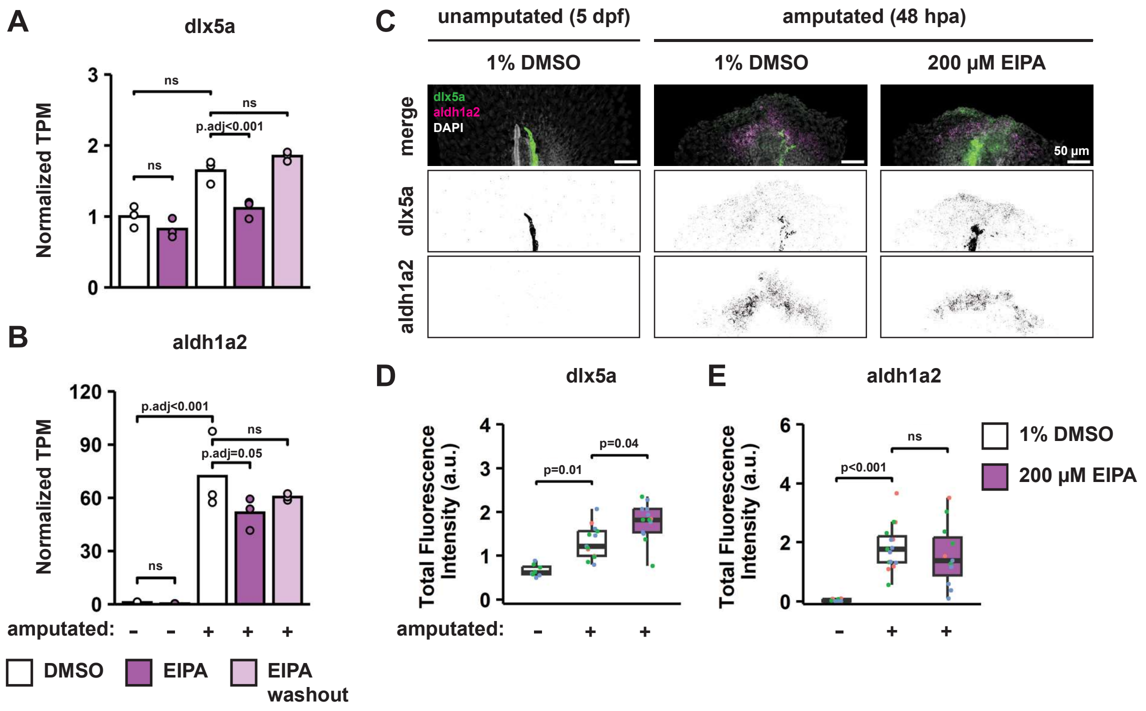


**Fig. S9 - Effect of EIPA treatment on wound epithelium and blastema markers**

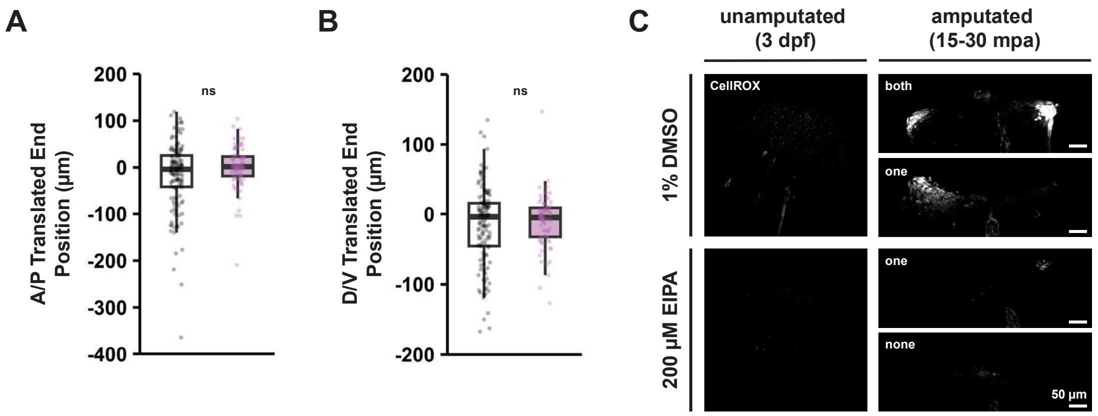


**Fig. S10 - Neutrophil end positions and single-channel CellROX images**

**Fig. S1 – mKate2 and SEpHluorin dynamics**

(A) Representative maximum projection images of unamputated and amputated *Tg(ef1α:mKate2-SEpHluorin)* larval tails at 3 dpf over 12 h. (B,C) SEpHluorin (B) and mKate2 (C) fluorescence intensity measured in amputated *Tg(ef1α:mKate2-SEpHluorin)* larval tails (0-40 µm from wound) incubated for 20 min in nigericin calibration buffers plotted vs. pH of buffers. Dots represent fish. Dot shapes represent experimental days. n=28.

**Fig. S2 – pHi dynamics post-amputation persists up to 120 µm from the wound site**

(A-D) Time course of SEpHluorin/mKate2 ratios measured in unamputated and amputated *Tg(ef1α:mKate2-SEpHluorin)* larval tails at indicated distances from wound normalized to t=0 timepoint for each fish. Error bars show SEM. n=5,5,3. p-values from Kruskal-Wallis and Wilcoxon signed rank tests (mu=1). Kruskal-Wallis p-values in unamputated groups: n.s. (A-D); 1% DMSO amputated groups: p=0.001 (A), p=0.02 (B), n.s. (C,D); 200 µM EIPA amputated groups: p<0.001 (A-D).

**Fig. S3 – Exposure to EIPA or low pHe has no effect on larval survival or tail growth**

(A,D) Survival curves from aggregated counts across three replicates (starting n=16-20 unamputated wildtype fish per group per replicate). (B,C) Representative images (B) and area (C) of unamputated wildtype larval tails in 1% DMSO or 200 µM EIPA at 6 dpf. p-value from t-test. (E,F) Representative images (E) and area (F) of unamputated wildtype larval tails in indicated pHe at 6 dpf. p-value from Kruskal-Wallis test. (C,F) Dots represent fish. Dot colors represent experimental days. Boxplots show median and interquartile range. n=15 per group.

**Fig. S4 – CRISPR/Cas9-mediated knockout of NHE family member genes**

(A) Phylogram of *Homo sapiens* (Hs) and *Danio rerio* (Dr) NHE family member protein sequences. (B) Area of regrowth measured in amputated crispant larval tails at 3 dpa. n=16-28 per group. p-values from one-way ANOVA (p<0.001) and Dunnett's test vs. scramble. (C) Area of unamputated crispant larval tails measured at 6 dpf. n=14-15 per group. p-value from one-way ANOVA. (D-G) Area of regrowth measured in amputated larval tails at 3 dpa. Fish were F2 progeny from incrosses between stable heterozygous carriers of indicated mutant alleles. n=5-21 per group. p-values from one-way ANOVA or Kruskal-Wallis tests (n.s.). (B-G) Dots represent fish. Dot colors represent experimental days. Boxplots show median and interquartile range.

**Fig. S5 – Bulk RNA-seq at 6 hpa identifies role for cholesterol biosynthesis during tail regeneration**

(A) Schematic of bulk RNA-seq conditions at 6 hpa. (B) Principal component analysis (PCA) plot of bulk RNA-seq results at 6 hpa. Dots represent samples. Dot shapes represent replicates. (C) Gene ontology (GO) terms enriched among differentially expressed genes between EIPA- and DMSO-treated larvae at 6 hpa. (D) Volcano plot of 6 hpa EIPA vs. DMSO comparison. Dots represent genes. Downregulated genes with adjusted p-value < 0.05 are shown in black (or orange to highlight cholesterol biosynthesis genes). Upregulated genes with adjusted p-value < 0.05 are shown in purple. (E) Schematic of cholesterol biosynthesis pathway with enzymes and pharmacological inhibitor highlighted in orange. (F,G) Representative images (F) and area of regrowth (G) of wildtype larval tails amputated in increasing concentrations of atorvastatin at 3 dpa. n=16-21 per group. p-values from one-way ANOVA (p<0.001) and Dunnett's tests. vs. 1% DMSO. (H) Survival curves from aggregated counts across three replicates (starting n=20 unamputated wildtype fish per group per replicate). (I,J) Representative images (I) and area (J) of unamputated wildtype larval tails in 1% DMSO or 10 µM atorvastatin at 6 dpf. n=15,14. p-value from t-test. (G,J) Dots represent fish. Dot colors represent experimental days. Boxplots show median and interquartile range.

**Fig. S6 – Summary of bulk RNA-seq at 48 hpa**

(A) Table of 48 hpa differentially expressed genes (DEGs) defined by adjusted p-value < 0.05 and $|\log_2$ fold change$| > 0.5$. (B) Venn diagram of 48 hpa DEGs comparing effects of full-length EIPA and EIPA washout treatments in amputated and unamputated larvae. (C) Venn diagram of 48 hpa upregulated and downregulated genes during regeneration in EIPA vs. DMSO. (D) Transcripts per million (TPM) (log transformed and z-score normalized by row) of cell cycle and pro-inflammatory genes from bulk RNA-seq samples at 48 hpa. Columns show individual replicates.

**Fig S7 – Single-channel phospho-H3 images and EdU labeling with EIPA**

(A,B) Representative maximum projection images of phospho-H3 immunolabeling in unamputated and amputated wildtype larval tails. Dotted lines indicate boundaries of ROIs used for quantification. (C,D) Representative maximum projection images (C) and quantifications (D) of EdU labeling in amputated wildtype larval tails at 48 hpa. Dotted lines indicate boundaries of ROIs used for quantification. Dots represent fish. Dot colors represent experimental days. Boxplots show median and interquartile range. n=19,17. p-value from Wilcoxon rank sum test.

**Fig S8 – Exposure to EIPA but not low pHe leads to increased cell death**

(A,B) Representative maximum projection images (A) and quantifications (B) of TUNEL labeling in unamputated and amputated wildtype larval tails in 1% DMSO or 200 µM EIPA. n=12-21 per group. p-values from Kruskal-Wallis (p<0.001) and Dunn's tests. (C,D) Representative maximum projection images (C) and quantifications (D) of TUNEL labeling in amputated wildtype larval tails in indicated pHe at 48 hpa. n=9,10. p-value from t-test. (B,D) Dots represent fish. Dot colors represent experimental days. Boxplots show median and interquartile range.

**Fig. S9 – Effect of EIPA treatment on wound epithelium and blastema markers**

(A,B) Normalized *dlx5a* (A) and *aldh1a2* (B) transcripts per million (TPM) counted from bulk RNA-seq of unamputated and amputated wildtype larval tails at 48 hpa. Adjusted p-values are from DESeq2 differential gene expression analysis. (C) Representative maximum projection images of *dlx5a* and *aldh1a2* RNA-FISH in unamputated and amputated wildtype larval tails. (D,E) *dlx5a* (D) and *aldh1a2* (E) total fluorescence intensity quantified in unamputated and amputated wildtype larval tails. Dots represent fish. Dot colors represent experimental days. Boxplots show median and interquartile range. n=9-15 (D) or n=8-13 (E) per group. p-values from Kruskal-Wallis ($p<0.001$) and Dunn's tests.

**Fig. S10 – Neutrophil end positions and single-channel CellROX images**

(A,B) End position at 4 hpa along the anterior/posterior (A/P) axis (A) or dorsal/ventral (D/V) axis (B) for the subset of neutrophils whose start position was close to the amputation plane in *Tg(mpx:dendra2); Tg(cdh1:cdh1-tdTomato)* larval tails. Tracks were translated to start at position (0,0). In (A), positive values indicate posterior movement, and negative values indicate anterior movement. In (B), positive values indicate dorsal movement, and negative values indicate ventral movement. Dots represent cells. Boxplots show median and interquartile range. n=131,74 cells. N=12 fish per group. p-values from Wilcoxon rank sum tests. (C) Representative maximum projection images of CellROX staining for ROS in unamputated and amputated wildtype larval tails.

**Table S1 – Differential gene expression analysis**

Results from DESeq2 differential gene expression analysis of different pairwise comparisons (indicated in sheet tab names) from bulk RNA-seq data.

**Table S2 – Biological process gene ontology enrichment analysis**

Results from biological process gene ontology enrichment tests conducted on different sets of differentially expressed genes (indicated in sheet tab names) between amputated EIPA- and DMSO-treated larvae.

**Movie S1 – Neutrophil dynamics in unamputated DMSO-treated larval tail**

Unamputated *Tg(mpx:dendra2); Tg(cdh1:cdh1-tdTomato)* larval tail in 1% DMSO showing neutrophils in orange and cell membranes in grey over 18 min. Scale bar: 50 µm.

**Movie S2 – Neutrophil dynamics in DMSO-treated larval tail from 0 to 4 hpa**

Amputated *Tg(mpx:dendra2); Tg(cdh1:cdh1-tdTomato)* larval tail in 1% DMSO showing neutrophils in orange and cell membranes in grey over 4 hpa. Scale bar: 50 µm.

**Movie S3 – Neutrophil dynamics in unamputated EIPA-treated larval tail**

Unamputated *Tg(mpx:dendra2); Tg(cdh1:cdh1-tdTomato)* larval tail in 200 µM EIPA showing neutrophils in orange and cell membranes in grey over 18 min. Scale bar: 50 µm.

**Movie S4 – Neutrophil dynamics in EIPA-treated larval tail from 0 to 4 hpa**

Amputated *Tg(mpx:dendra2); Tg(cdh1:cdh1-tdTomato)* larval tail in 200 µM EIPA showing neutrophils in orange and cell membranes in grey over 4 hpa. Scale bar: 50 µm.
